# Ultrafast and long-range coordination of wound responses is essential for whole-body regeneration

**DOI:** 10.1101/2023.03.15.532844

**Authors:** Yuhang Fan, Chew Chai, Pengyang Li, Xinzhi Zou, James E. Ferrell, Bo Wang

**Affiliations:** Department of Bioengineering, Stanford University, Stanford, CA, USA; Department of Chemical and Systems Biology, Stanford University School of Medicine, Stanford, CA, USA; Department of Biochemistry, Stanford University School of Medicine, Stanford, CA, USA

## Abstract

Injury induces systemic, global responses whose functions remain elusive. In addition, mechanisms that rapidly synchronize wound responses through long distances across the organismal scale are mostly unknown. Using planarians, which have extreme regenerative ability, we report that injury induces Erk activity to travel in a wave-like manner at an unexpected speed (∼1 mm/h), 10-100 times faster than those measured in other multicellular tissues. This ultrafast signal propagation requires longitudinal body-wall muscles, elongated cells forming dense parallel tracks running the length of the organism. Combining experiments and computational models, we show that the morphological properties of muscles allow them to minimize the number of slow intercellular signaling steps and act as bidirectional superhighways for propagating wound signals and instructing responses in other cell types. Inhibiting Erk propagation prevents cells distant to the wound from responding and blocks regeneration, which can be rescued by a second injury to distal tissues within a narrow time window after the first injury. These results suggest that rapid responses in uninjured tissues far from wounds are essential for regeneration. Our findings provide a mechanism for long-range signal propagation in large and complex tissues to coordinate cellular responses across diverse cell types, and highlights the function of feedback between spatially separated tissues during whole-body regeneration.

## Introduction

It has long been noted that injury can induce responses in uninjured tissues far from wounds (Losner et al., 2021). In many invertebrates, injury triggers proliferation not only in cells nearby but also in those millimeters to centimeters away (Ricci and Srivastava, 2018; Wenemoser et al., 2012), suggesting wound signals can spread across long distances even though these organisms lack a circulatory system. In mouse, injury to muscles in one leg causes stem cells in the contralateral leg to switch from quiescence to an alert state, which may prepare tissues for future injuries (Rodgers et al., 2014; Rodgers et al., 2017). Similar effects have been observed after axolotl limb amputation, implicating that long-range induction of wound responses may be a broadly conserved phenomenon (Johnson et al., 2018; Payzin-Dogru et al., 2021). More recently, cardiac injury in zebrafish was shown to induce coordinated gene expression changes in distant organs including brain and kidney controlled by a single permissive enhancer, but attempts to eliminate the distal responses did not alter the heart regeneration outcome (Sun et al., 2022). Thus, although systemic wound responses appear to be widespread, it remains unclear whether they contribute to the current round of regeneration or simply represent byproducts of injury-induced signaling cascades.

In order to possibly participate in regeneration, the distal wound responses need to turn on shortly after injury within the right time window as the regeneration proceeds. This time scale should be determined by the rate at which wound signals are communicated between injury and distant sites. However, besides circulatory factors (Halme et al., 2010; Rodgers et al., 2017; Hirose et al., 2019), mechanisms that can rapidly transmit molecular signals over millimeter-to-centimeter distances in nonvascularized tissues are largely unknown.

While wound signals may spread in the form of diffusive cues (Chera et al., 2009; Hasegawa et al., 2015; Kikuchi et al., 2011), a long-standing puzzle is that diffusion is often too short-ranged in densely packed tissues (Müller et al., 2013). It has been recently shown that coupling diffusion with biochemical positive feedbacks can induce travelling waves and help to overcome some limitations of simple diffusion (Aoki et al., 2017; Hiratsuka et al., 2015; De Simone et al., 2021). However, due to slow intercellular communication in multicellular tissues, the observed propagation speeds of these waves, ∼10-100 µm/h (Hayden et al., 2021; De Simone et al., 2021), are still incompatible with the fast, long-range communication required by regeneration programs that need to rapidly progresses in time.

To study the coordination and function of systemic wound responses, we investigate the planarian flatworm *Schmidtea mediterranea*, as it has a remarkable ability to regenerate essentially any missing body parts (Newmark and Sánchez Alvarado, 2002; Rink, 2013; Reddien, 2018). They can regrow into normal healthy organisms from minute tissue remnants on a time scale of days. Injury induces broad transcriptional changes throughout the animal (Wenemoser et al., 2012). Accompanying these global molecular responses is elevated stem cell proliferation (Wenemoser and Reddien, 2010), which is then followed by the induction of transient regeneration-activated cell states in various tissue types (Benham-Pyle et al., 2021) to reestablish body polarity (Scimone et al., 2017), sustain stem cell proliferation, and initiate tissue remodeling.

We sought to understand how this long-range coordination arises and whether the wound responses in uninjured tissues are required for regeneration. We found the wound signal to propagate in the form of an Erk wave that travels at an unexpected speed (∼1 mm/h), 10-100 times faster than those reported in other multicellular tissues (Hiratsuka et al., 2015; De Simone et al., 2021). The ultrafast propagation of Erk activity is enabled by the longitudinal body-wall muscles, which act as superhighways for signal transduction and relay the wound signal to other cell types instructing responses therein. Combining experiments and a theoretical model, we proposed that the morphological properties of muscle cells, i.e., close packing of parallel elongated cell bodies, provide the cellular basis for rapid long-range communication. Inhibiting Erk activity propagation and thereby distal wound responses blocked planarian regeneration, which was rescued by inducing wound responses through a second amputation of distal tissues within a narrow time window after the first injury. These findings suggest that proximal responses alone are insufficient to drive regeneration and timely long-range feedback between spatially separated tissues is essential for whole-body regeneration.

## Results

### Rapid spatial coordination of the planarian wound responses is mediated by ultrafast Erk activity waves

To better characterize the spatial coordination of planarian wound responses at the systems level, we performed RNAseq on tails after amputating tail and head regions respectively (Figure 1A). This experiment allowed us to quantitatively compare responses proximal and distal to wounds within the same tissue type. Although the proximal responses were more pronounced initially, we observed a strong correlation between the proximal and distal responses at 6 hour post amputation (hpa) (Figure 1B-C and S1A), except for a focal set of genes specifically induced at the wound (Figure 1C). Such global correlation was maintained at 24 hpa even though the upregulated genes were different (Figure S1B-C).

**Figure 1.**
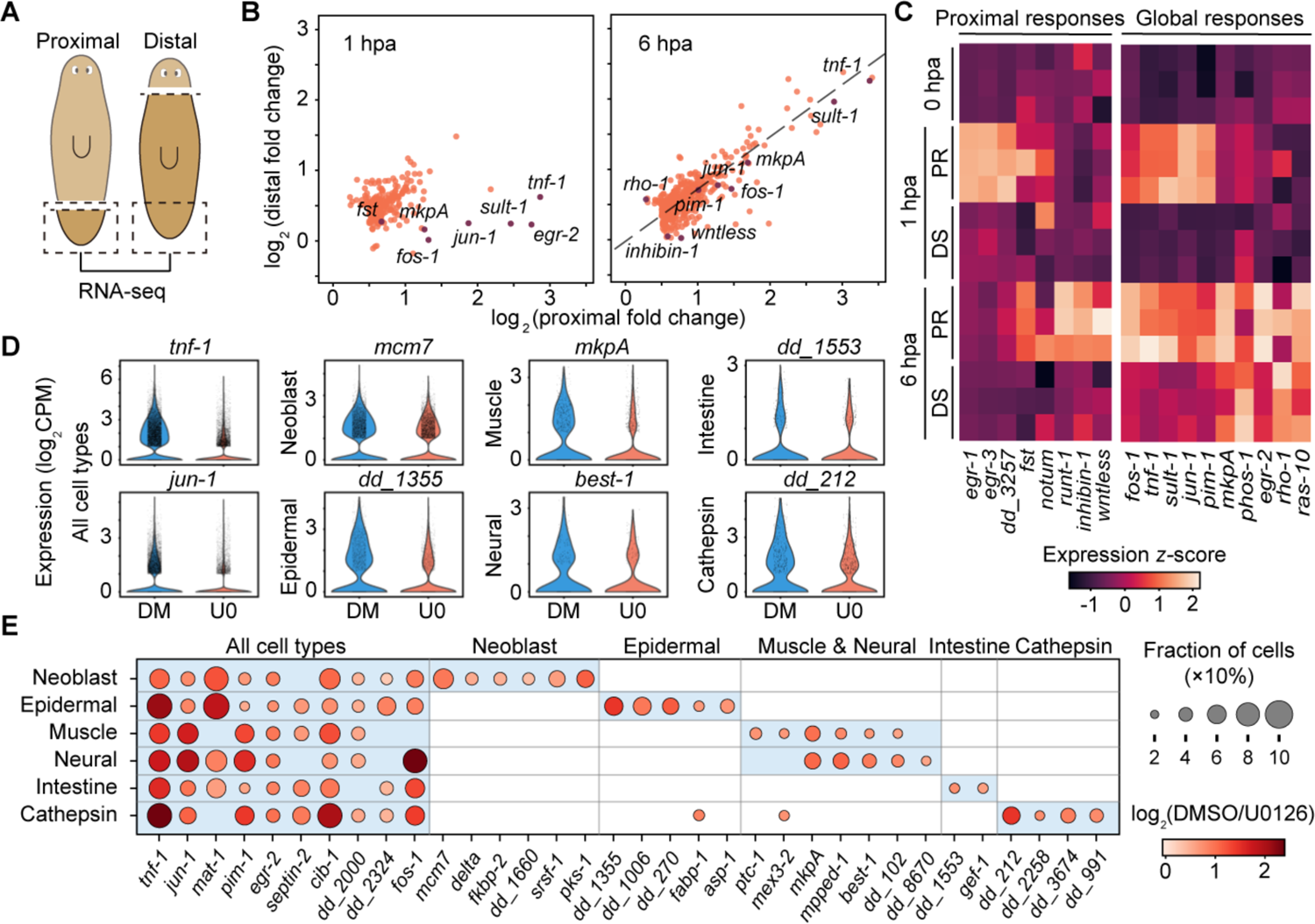
Coordination of planarian wound responses across long distances and broad cell types. (A) Strategy to measure proximal and distal wound responses in the same tissue. (B) Comparison of gene upregulation proximal and distal to wounds. Genes plotted have p-value<0.001 (two-sided Welch’s t-test) and log_2_(fold change)>0.5 in either proximal or distal samples, each containing three biological replicates. (C) Heatmap of example upregulated genes. PR, proximal; DS, distal. Global responses include genes activated in both proximal and distal responses. (D) Violin plots showing expression distribution of wound response genes in animals treated by DMSO (DM) or 25 μM U0126 (U0). Points: data of individual cells. Genes shown are downregulated in U0126 treated samples (p-value<0.01, two-sided Mann-Whitney-Wilcoxon test) in cell types specified. Data in all cell types are provided in Figure S3. (E) Expression fold changes of Erk-dependent global wound response genes in individual cell types. Genes shown are upregulated at 6 hpa globally measured by bulk RNAseq (p-value<0.001, Two-sided Welch’s t-test, Figure S1D) and downregulated by U0126 treatment (p-value<0.01, two-sided Mann-Whitney-Wilcoxon test) in specific cell types according to the scRNAseq data.

Planarian wound responses depend upon the activation of extracellular signal-regulated kinase (Erk) (Figure S1D-E) (Owlarn et al., 2017; Tasaki et al., 2011), via a highly conserved wound repair pathway (Chera et al., 2009; DuBuc et al., 2014; Mace et al., 2005; Srivastava, 2021; Tursch et al., 2020). This raises the possibility that Erk signaling is responsible for coordinating planarian wound responses. Indeed, in mouse skin (Hiratsuka et al., 2015) and zebrafish scales (De Simone et al., 2021), Erk activity can be relayed between cells in the form of trigger waves after injury (Figure S1F). Trigger waves are self-regenerating fronts of activity that can be produced in signaling systems with positive feedback, and they spread without losing speed or amplitude, making them suitable for long range signaling (Deneke and Di Talia, 2018; Gelens et al., 2014; Tyson and Keener, 1988; Winfree, 1974). However, the reported Erk wave speeds (∼10-100 µm/h) are orders of magnitude too slow to explain the rapid activation of distal wound responses in the planarian. It would take days or even weeks for the signal to travel across the planarian body, which is typically 5-20 mm long.

In addition, previous studies of Erk waves in multicellular systems have focused on tissues comprising uniform cell populations (Aoki et al., 2017; Gagliardi et al., 2021; Hayden et al., 2021; Hiratsuka et al., 2015; De Simone et al., 2021). In contrast, planarian wound responses need to be coordinated across various cell types that may have different sensing mechanisms, activation kinetics, and competency in transmitting the signal. Indeed, using single-cell RNAseq (scRNAseq) on tissues at 3 hpa in the presence or absence of the Erk kinase (Mek) inhibitor U0126 (Figure S2), we found that, along with cell-type specific responses (e.g. activation of *mcm7* in neoblasts, *mkpA* in muscles and neurons), generic wound responses (e.g., activation of *tnf-1*, *jun-1*) were induced in most cells after injury, in an Erk-dependent manner (Figure 1D-E and S3).

These reservations notwithstanding, we examined whether Erk signal propagates in a wave-like manner in planarian tissues after injury. We quantified Erk activity (defined by the ratio between phosphorylated Erk, pErk, and total Erk) in tissue pieces evenly spaced ∼1 mm apart using western blotting (Figure 2A and S4A-B). In line with the timing of the transcriptional changes (Figure 1B-C), Erk activity peaked around 0.5 hpa in tissues proximal to wound, and propagated distally with ∼1 h delay in peak time between adjacent positions (Figure 2A-B). The linear relationship between peak time and distance from wounds is consistent with wave propagation. In addition, the signal amplitude did not decrease with distance travelled (Figure 2A-B). This can account for the high correlation between proximal and distal transcriptional wound responses (Figure 1B).

**Figure 2.**
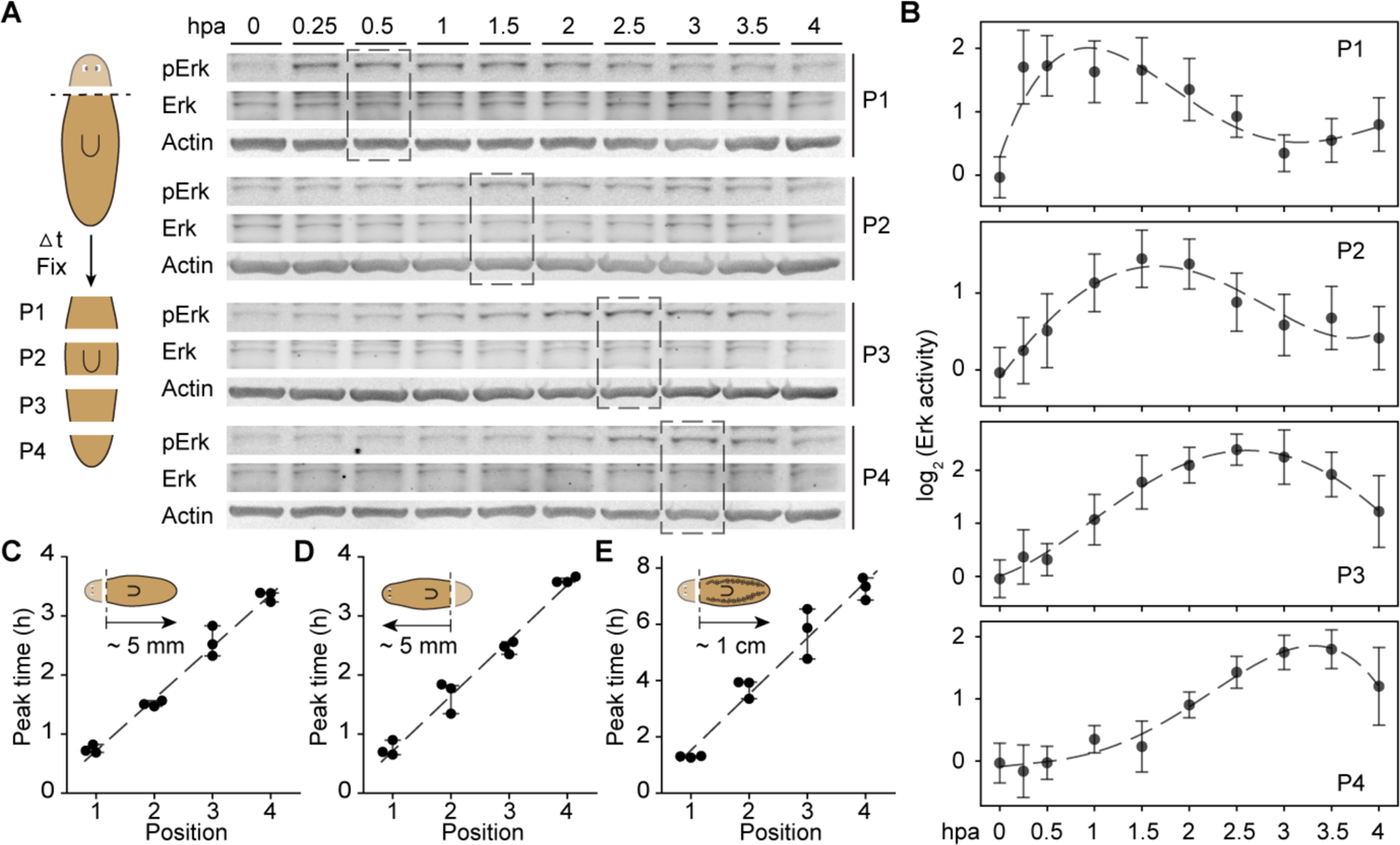
Ultrafast propagation of Erk activity after injury. (A) Schematic and representative images of the western blotting experiments. Loading control: actin. Dashed boxes, maximal Erk activity. (B) Quantification of Erk activity from western blot using the ratio between pErk and total Erk intensities. Dashed lines: polynomial fit. (C-E) Peak times vs. proximal-to-distal position. Dashed line: linear fit. Insets: amputation planes and length of animals used in experiments. Planarians of the asexual biotype were used in (C, D), and the sexual biotype in (E) as they have longer bodies. Error bars: standard derivation (SD). Raw data for (D-E) are shown in Figure S4C.

From the time and distance relationship, we measured the wave speed to be ∼1 mm/h (Figure 2C), which is 1-2 orders of magnitude faster than reported values in other multicellular tissues (Aoki et al., 2017; Hiratsuka et al., 2015; De Simone et al., 2021) but comparable to the speed of some intracellular trigger waves in large cells (Chang and Ferrell, 2013; Cheng and Ferrell, 2018). This speed was independent of propagation direction—that is, Erk activity could spread either from head to tail or from tail to head, depending on where the wound was located (Figure 2D and S4C)—and was independent of animal size (Figure 2E and S4D). These findings suggest that wave speed is controlled by the intrinsic properties of a bidirectional signaling system.

### Distal wound responses are essential for regeneration

One prediction of trigger wave models is that the speed of the Erk wave should be reduced by partially inhibiting the intracellular Erk cascade (Gelens et al., 2014; Hayden et al., 2021). Consistent with this prediction, Erk activation in distal tissues was significantly delayed with increasing Mek inhibitor (U0126) concentrations and was eventually blocked when the U0126 concentration exceeded a critical threshold (∼8 μM), whereas Erk activity was only slightly reduced at wounds through a broad range of inhibitor concentrations (Figure 3A-B). This allowed us to evaluate regeneration phenotypes while tuning the rate of Erk activity propagation and to separate the contributions of proximal and distal wound responses during regeneration.

**Figure 3.**
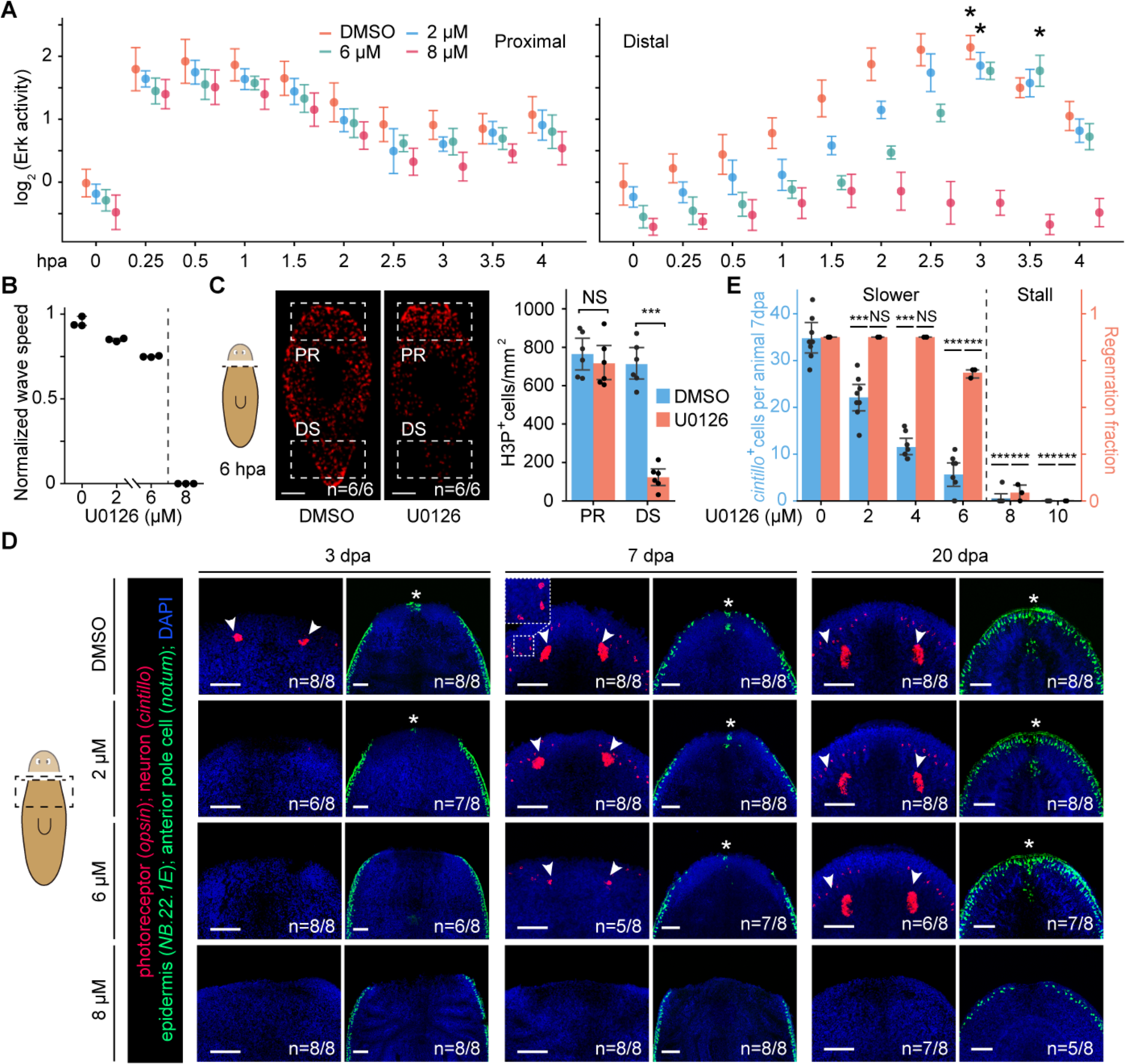
Erk activity propagation is essential to planarian regeneration. (A) Dependence of Erk activity, in tissues proximal (P1, left) and distal (P3, right) to wounds, on U0126 concentration. Asterisks: peak activities in distal responses. (B) Fitted wave speed vs. U0126 concentration. (C) (Left) Anti-H3P labels mitotic cells at 6 hpa in animals treated with DMSO or 8 μM U0126. (Right) Number of H3P^+^ cells per area in proximal (PR) and distal (DS) regions (boxes in images). (D) Regeneration of anterior tissues is delayed by increasing concentrations of U0126. Asterisks: *notum* expression at anterior pole; arrows: *opsin* expression in photoreceptors. Inset: *cintillo*^+^ neurons. (E) Number of regenerated *cintillo*^+^ neurons (eight animals per condition), and fraction of regenerated animals (three replicates each containing twenty animals) vs. U0126 concentration. In schematics, dashed lines: amputation planes; box: imaging area. ∗∗∗, p < 0.001; NS, no significant difference, two-sided Welch’s t-test; error bars: standard derivation (SD) in (A-B), 95% confidence interval (CI) in (C) and (E); n: number of samples consistent with the image out of the total number of samples analyzed. Scale bars, 500 μm in (C), 100 μm in (D).

We found that blocking Erk propagation inhibited stem cell mitosis only in distal tissues post amputation, but left proliferation at wound sites unaffected (Figure 3C and S4E). This result establishes a direct link between Erk activation and stem cell proliferation. The regeneration of anterior tissues (e.g., *opsin*^+^ photoreceptors and *notum*^+^ anterior pole cells) after decapitation was also delayed with slower Erk propagation and absent when Erk activation was blocked in distal tissues, although the epidermis (*NB.22.1E*^+^) eventually closed over wounds at a much later time (Figure 3D). To quantify the rate of regeneration, we counted *cintillo*^+^ chemosensory neurons regenerated at 7 dpa and found that their numbers decreased with reduced Erk propagation until dropping to zero in animals treated with U0126 above the critical concentration (Figure 3E).

We noticed that Erk activity propagation was most critical to regeneration when wound responses needed to be activated across long distances. When we sliced planarians such that the majority of the remaining tissues had been exposed to injury, regeneration was less sensitive to U0126 treatment, as the proximal wound responses were mostly unaffected around the critical inhibitor concentration (Figure 4A).

**Figure 4.**
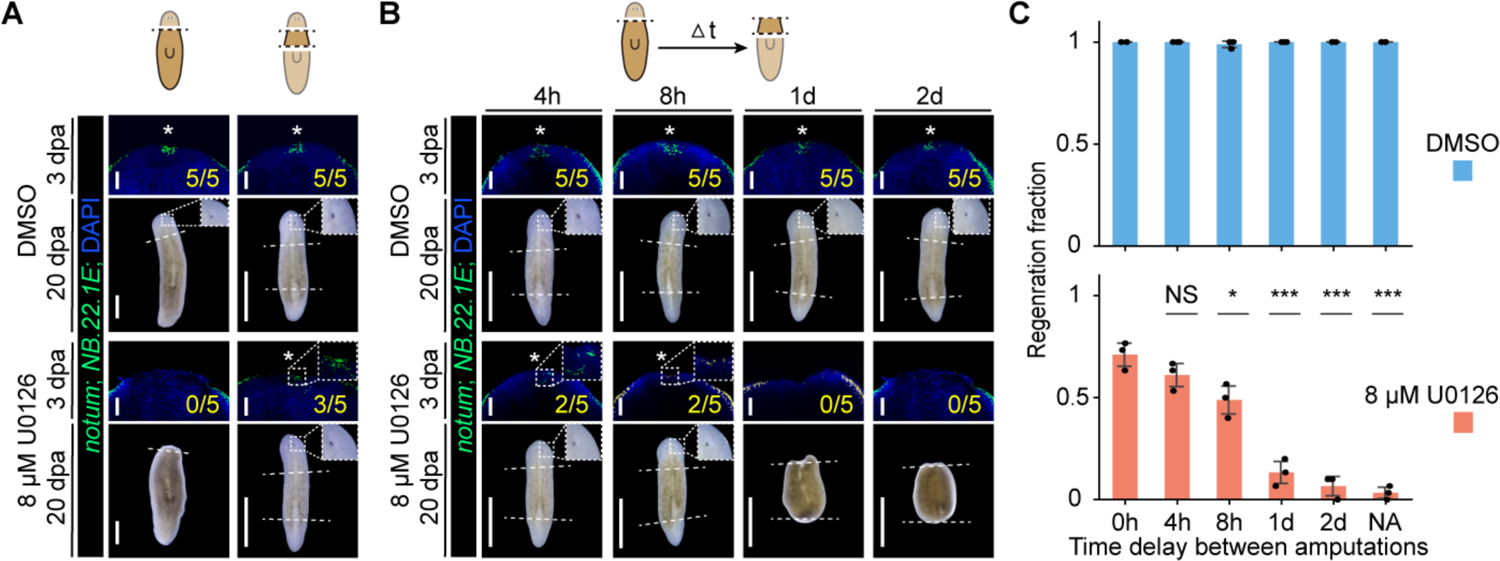
Distal wound responses are essential for planarian regeneration. (A) Anterior regeneration is fully blocked by 8 μM U0126 treatment after single amputation but only weakly affected after double amputation. At this U0126 concentration, distal responses are eliminated, but proximal wound responses are mostly unaffected. (B) Second amputation within a few hours of the first amputation is sufficient to rescue regeneration deficiencies caused by 8 μM U0126 treatment. In contrast, animals fail to regenerate head or tail if there is a long time delay between the first and second amputations. In (A-B), scale bars: 100 μm in FISH images, 1 mm in bright field images. Dashed lines: amputation planes. Asterisks: *notum* expression at anterior pole. Insets: re-specified anterior pole cells in FISH images and regenerated photoreceptors imaged on live animals. Animals treated with DMSO are used as controls and can fully regenerate regardless of types of amputation. (C) Fraction of regenerated animals (three replicates each containing thirty animals) treated by DMSO or 8 μM U0126 vs. time delays between the first and second amputations. NA: no second amputation. ∗, p < 0.05; ∗∗∗, p < 0.001; NS, no significant difference, two-sided Welch’s t-test, compared against the 0 h data; error bars: standard derivation (SD).

Motivated by this surprising result and given that distal and proximal wound responses are largely correlated, we reasoned that inducing responses in distal tissues through a second injury may compensate for the lack of long-range coordination of wound responses. Whereas the delay time between proximal and distal responses under natural conditions should be defined by the speed at which wound signals propagate, using two amputations separated in time (Figure 4B), we can tune the delay between the first and second injury to investigate the effects of temporal coordination of wound responses.

To test this hypothesis, we decapitated the planarians treated with U0126 at 8 μM to eliminate the distal wound responses, waited for various amounts of time, and then amputated the tail. Strikingly, a second amputation within a few hours of the first amputation was sufficient to rescue regeneration, whereas the planarians with tails amputated a day after the first amputation failed to regenerate either head or tail (Figure 4B-C). The regeneration was stalled at an early stage even before the reset of body polarity, manifested by the lack of both *notum* expression and blastema formation. Surprisingly, not only did the first amputation in heads require the second tail injury to regenerate, but tail regeneration from the second amputation also relied on a recent first head injury. Altogether, our findings demonstrate that wound responses in tissues distant to wounds depend on the propagation of Erk activity; delayed or missing distal Erk activation can cause regeneration deficiencies, stressing the necessity of ultrafast propagation of Erk activity.

### Longitudinal body-wall muscles are required for Erk activity propagation and the activation of distal wound responses

The observations above raise the question of how Erk activation can propagate so quickly. Previous analyses have shown that the relay of Erk activation between cells is the rate limiting step (Figure S1F) (Hayden et al., 2021; De Simone et al., 2021), whereas intracellular signaling is relatively fast and can reach maximal response within minutes (Kiyatkin et al., 2020). Therefore, we hypothesized that reducing the number of intercellular events per unit distance by increasing the size of cells that relay Erk signal might significantly speed up wave propagation. We formalized this hypothesis using a diffusive signaling relay model (Dieterle et al., 2020). Our simulation indicated that when intracellular propagation is fast relative to intercellular propagation, wave speed increases with propagating cell size (Figure S5A-D). This analysis suggested that cell types with long cell bodies could act as superhighways for signal propagation in heterogeneous tissues containing cells of a broad range of sizes.

While most planarian cells are small (∼5-10 µm diameter), the body wall muscles are mononucleate cells over 100 µm long and assemble into a dense, continuous network covering the whole animal body, with sub-micron spacing between muscle fibers (Scimone et al., 2017; Witchley et al., 2013; Lim et al., 2019) (Figure 5A). We found that Erk was phosphorylated in most muscle cells after injury (Figure 5A and S6A-B). Ablation of the longitudinal muscle fibers through RNAi-mediated silencing of *myoD*, a muscle-specific transcription factor required for longitudinal muscle specification in the planarian (Scimone et al., 2017), led to a marked reduction of Erk activation at wounds, which is consistent with the idea that longitudinal muscles are the main early responders to injury. *myoD* knockdown also fully blocked injury-induced Erk activation in distal tissues, lending support to our hypothesis (Figure 5B and S6C). Concordantly, >80% of the global wound response genes failed to respond to injury in distal tissues after *myoD* RNAi, though they maintained their responses at proximal sites (Figure 5C-D). As the *myoD*-dependent wound response genes had significant overlap with Erk-dependent genes (Figure 5E), our results suggest that *myoD* RNAi affects wound responses mainly through Erk signaling.

**Figure 5.**
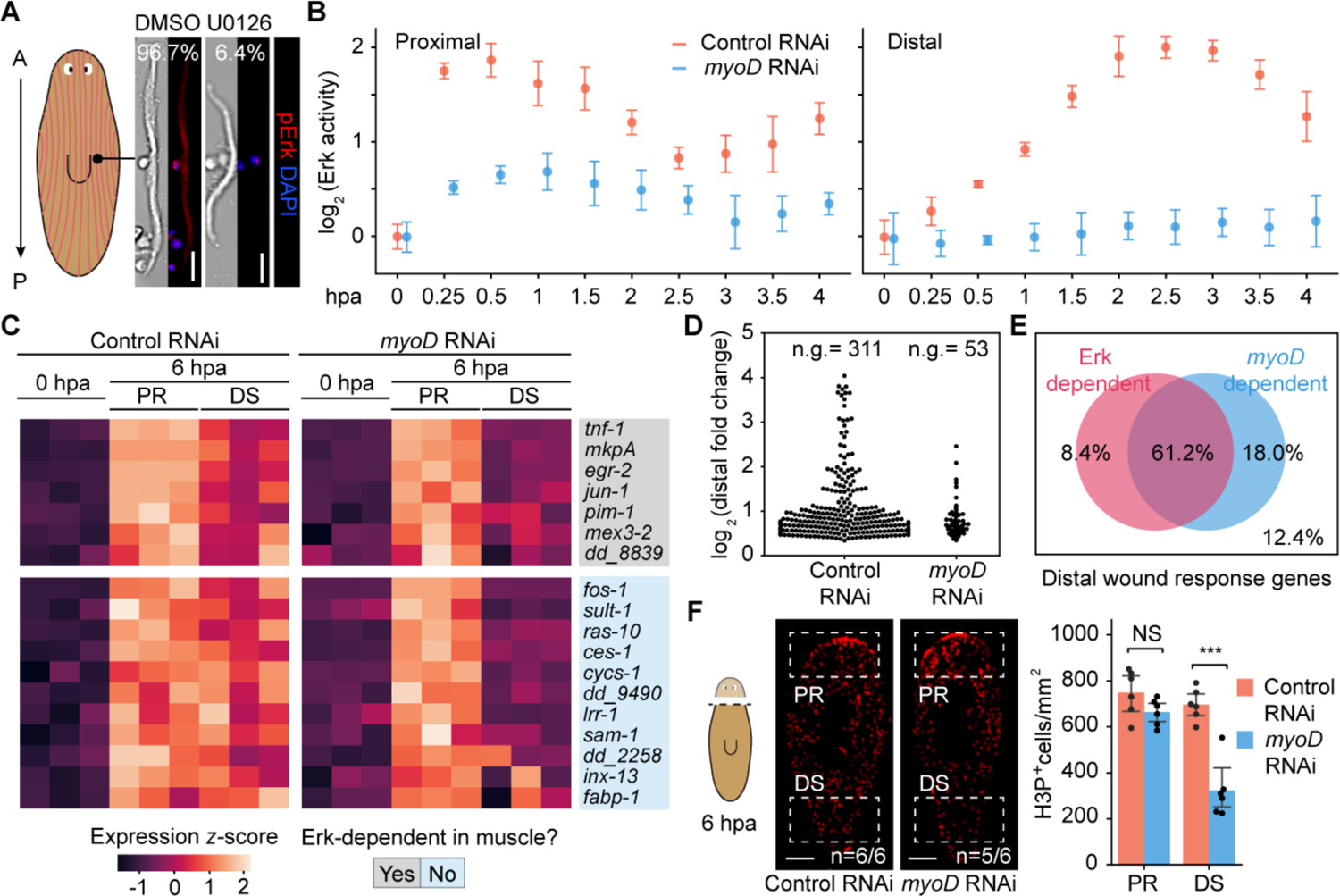
Longitudinal body-wall muscles are required for Erk activity propagation and distal wound responses. (A) (Left) Schematic of planarian longitudinal body-wall muscles (red lines). (Right) Bright-field and pErk immunofluorescence images of muscle cells isolated from wounded planarians (at 3hpa) treated with DMSO or 25 μM U0126. %, pErk^+^ fraction of muscle cells. (B) Erk activation in proximal tissues is reduced by *myoD* RNAi, and completely blocked in distal tissues. The Erk activity is determined by western blotting (Figure S6C). (C) Heatmap showing *myoD* RNAi blocks gene activation in distal (DS) but not proximal (PR) tissues. Upregulation of these genes after injury also requires Erk activation (Figure S1D). Erk-dependence in muscle cells was determined using the scRNAseq data. (D) Fold change of wound responses at 6 hpa (p-value<0.001; Two-sided Welch’s t-test, compared to 0 hpa) in distal tissues; n.g., number of genes. (E) Overlap of Erk-dependent and *myoD*-dependent wound response genes in distal tissues. (F) (Left) Representative images of anti-H3P staining showing cell proliferation was reduced only in distal tissues after *myoD* RNAi; n: number of samples consistent with the image out of the total number of samples analyzed. (Right) Number of H3P^+^ cells counted in proximal (PR) and distal (DS) regions (boxes in images). ∗∗∗, p < 0.001; NS: no significant difference, Two-sided Welch’s t-test; error bars: standard derivation (SD) in (B), 95% confidence interval (CI) in (F). Scale bars: 20 μm in (A), 500 μm in (F).

Importantly, within the overlap between Erk and *myoD*-dependent wound response genes, many of them were activated only in cell types other than muscles. Examples include *fos-1* in neurons and *ces-1* in neoblasts, as revealed by the scRNAseq analysis (Figure 5C and S1D). This observation highlights the non-autonomous role of muscle cells, which may function as ‘pioneers’ in transmitting the wound signal and disseminating it to other cell types. At the functional level, stem cell mitotic response in distal tissues was significantly reduced after *myoD* knockdown, whereas proliferation at the wound site was unchanged (Figure 5F), indicating that stem cell activation is Erk dependent and requires instructions from the muscle cells. While the planarian longitudinal muscles are already known to provide polarity cues for body plan reset during regeneration (Scimone et al., 2017; Witchley et al., 2013), our data reveal another essential function of the planarian muscles in facilitating the propagation of Erk waves and thereby coordinating wound responses across space and cell types.

### A diffusive signaling relay model identifies key cell morphological properties required to accelerate signal propagation

We extended the signaling relay model (Figure S5A-D) (Dieterle et al., 2020) to investigate morphological properties that may grant muscle cells special capacities in propagating Erk activity. Our model considered heterogeneous tissues containing both long cells, the major cell type that propagates the signal, and small round cells. We systematically tuned the length, volume density, and orientation of long cells, as well as the fraction of small cells that also can participate in the signal relay, in order to determine their effects on the speed of Erk activity wave (Figure S5E).

Through simulation, we found that the wave speed increased with increasing length of the long cells (Figure 6A and S5E). This trend was robust to changes of other model parameters such as long cell density and molecular kinetics of Erk signaling (Figure S5F-G). The wave speed was independent of long cell volume density in the low density regime, but above the critical density (∼0.2) where the long cells begin to form large continuous clusters (Figure S5H-I), the wave speed increased quickly with long cell density (Figure 6B). At the transition between these two density regimes, the size distributions of the clusters had a universal power-law shape (Figure 6C), which is reminiscent of ‘percolation’ transitions broadly observed in physical and biological systems (Stauffer and Aharony, 1994; Khariton et al., 2020). The nature of this transition implies that signal transmission via continuous paths formed by long cells connecting to each other and clustering in space is the major mode of the wave propagation. In addition, as the number of cells in alignment with the direction of signal propagation was increased, the wave propagation speed increased (Figure 6D and S5J). In contrast, the wave speed was only weakly dependent on the fraction of small cells that also can participate in the signal relay when the long cell density is low, and this dependence diminished quickly with increasing long cell density (Figure 6E and S5K).

**Figure 6.**
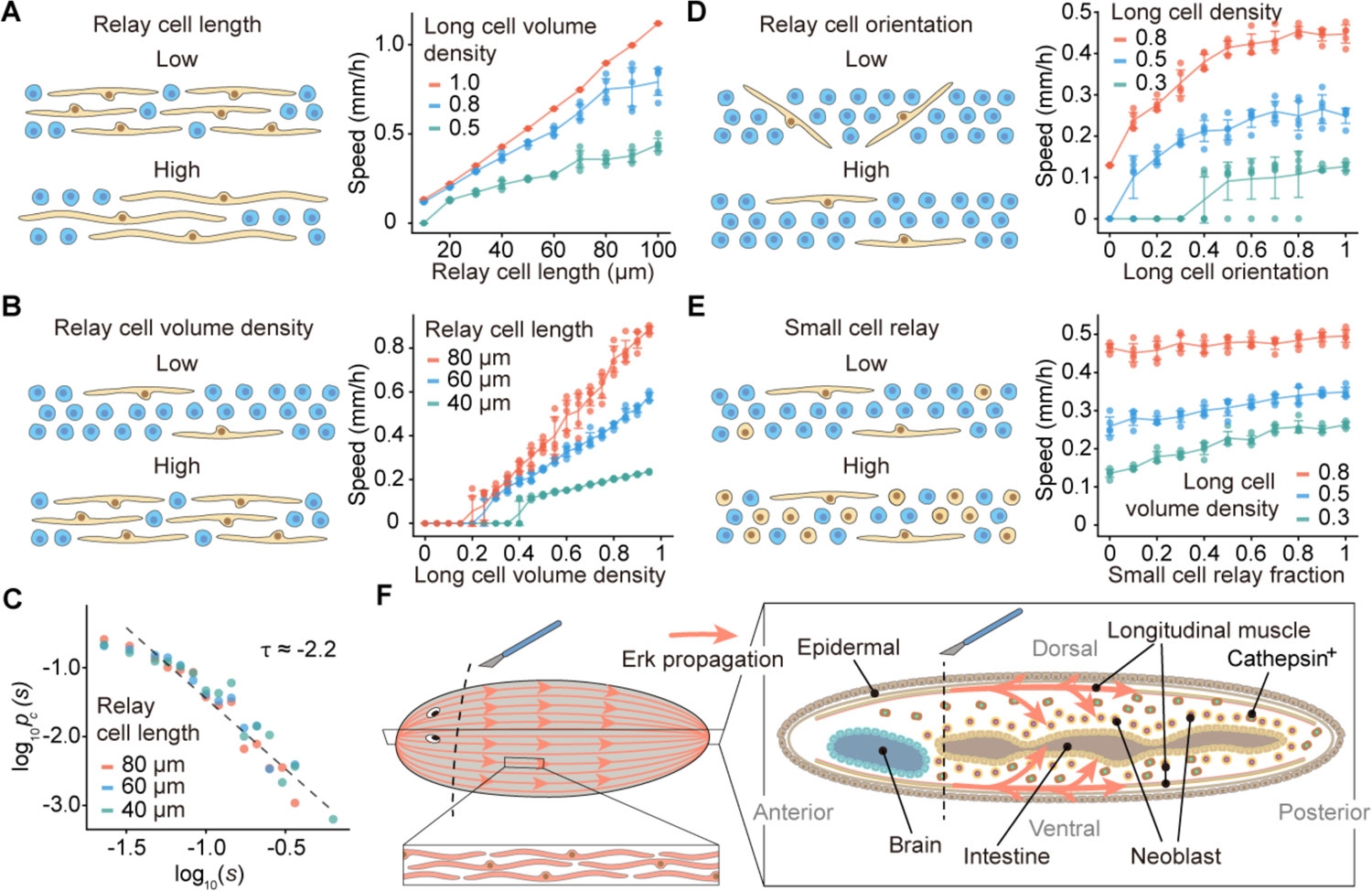
A diffusive relay model identifies cell morphological properties influencing signal propagation in heterogeneous tissues. (A) The speed of signal propagation increases with the length of relay cells. The number density is adjusted to match volume density for cells with different lengths. (B) Signal propagation speed increases with long cell volume density. The speed is infinitesimal below the percolation transition (see Figure S5H), which shifts towards smaller long cell density with increasing cell length. (C) Length distributions of long cell clusters near percolation transition exhibit universal power-law scaling. The dashed line, a guide for the eye, has an exponent of −2.2. (D) Signal propagation speed increases with the orientation factor of long cells, which is defined by the fraction of long cells extending along the wave direction whereas other long cells are perpendicular. (E) Signal propagation speed is weakly dependent on the fraction of small cells that can relay the signal. Except for simulations in (D), long cells are assumed to be aligned with the wave propagation direction. In (A-D), small cells can only receive signal. In (D-E), long cells are 50 μm in length. In schematics, yellow: relay cells, blue: receiving cells. (F) Model of the muscle cell function in propagating wound signals through Erk wave. Arrows: Erk signal.

Whereas previous modeling efforts mostly focus on the effects of varying kinetics that control activation within individual cells and communication between adjacent cells (Yde et al., 2011; Dieterle et al., 2020; Hayden et al., 2021), our model reveals the contribution of large scale structures formed by clusters of cells that collectively enhance the rate of signal propagation. Conceptually, this simple model conveys three important messages. First, in order to function as signaling superhighways, the propagating cells not only need to be long but also should form dense parallel tracks, which are exactly the obvious attributes of the planarian longitudinal body-wall muscles (Scimone et al., 2017). Second, in the presence of a superhighway cell type, other smaller cells may only have negligible contributions in propagating the signal but instead are activated by receiving the signal from long cells. Strictly speaking, these small cells are activated in a ‘phase wave’ (Winfree, 1974), which is distinct from but driven by the trigger wave travelling through the long cells that directly propagate the signal. This mixed mode of propagation may be a signature of signaling in heterogeneous tissues. Finally, as our model has no architecture specific to the planarian tissues, its major conclusions may be generalizable to other complex and heterogeneous multicellular systems to predict the effects of cell morphology and cell type diversity in determining signal propagation dynamics.

## Discussion

In this study, we revealed a mechanism for wound signals to traverse distances of several millimeters within just hours. We proposed the notion of ‘pioneer’ cells—cells that lead in signal transduction (at least over long distances) and then relay the signal to other cell types. Integrating experiments and computational modeling, we argued that these pioneer cells need to have special morphological properties in order to act as superhighways for rapid signal propagation in heterogeneous tissues. The data presented indicate that, in planarians, the longitudinal body-wall muscles play this role and are required to propagate the wound signal in the form of Erk waves, and to activate responses in tissues distant to wounds (Figure 6F). To draw an analogy, these muscle cells function much like a circulatory system for disseminating wound signals across long distances (Rodgers et al., 2014; Rodgers et al., 2017); however, no fluid flow is involved; instead, the biochemical signals propagate as a trigger wave through a diffusion-reaction based mechanism (Deneke and Di Talia, 2018; Gelens et al., 2014; Tyson and Keener, 1988; Winfree, 1974). The wave speed is unexpectedly fast for a multicellular system and roughly matches some intracellular waves in large cells (Chang and Ferrell, 2013; Cheng and Ferrell, 2018). In a way, the planarians respond to injury as rapidly as they would if they were gigantic unicellular organisms.

By perturbing Erk activity propagation, we provided, to the best of our knowledge, the first evidence in any animal that suggests distal wound responses are necessary for the current round of regeneration. In addition, in animals with Erk signal propagation blocked, amputating the distal tissues within the first few hours after the first injury is sufficient to rescue the regeneration deficiencies. Collectively, these observations reveal that the proximal and distal wound responses should have distinct functions and must interact with each other to enable regeneration.

At proximal sites, the injury-induced upregulation of a few key regulators previously known to be required for the planarian regeneration, including *follistatin* (*fst*) (Gaviño et al., 2013; Roberts-Galbraith and Newmark, 2013; Tewari, et al. 2018), *notum* (Petersen and Reddien, 2011), and *runt-1* (Wenemoser et al., 2012), are indeed Erk-dependent (Owlarn et al., 2017), but their activation is not observed in distal tissues. Intriguingly, the injury-induced expression of *fst* and *notum* has been noted to be mostly in *myoD*^+^ muscles facing wounds (Scimone et al., 2017), implicating a direct regulation of *fst* and *notum* transcription through the Erk pathway as a part of the proximal wound responses. Apparently, the function of distal wound responses is independent of any of these known regulators, but still appears essential to initiate the regeneration program, because regeneration stalls without distal wound responses before the re-specification of anterior pole cells, an early event during regeneration. Although we showed that neoblast proliferation in distal tissues requires Erk activation, it is unlikely that regeneration relies on these cells as the source of new tissues. Removing the distal tissues, and the neoblasts therein, does not inhibit regeneration. Consistently, neoblast proliferation is sustained at wounds for up to a week during the planarian regeneration but only transiently elevated in distal tissues (Wenemoser and Reddien, 2010). Planarian regeneration also involves apoptosis (Pellettieri et al., 2010) and remodeling of preexisting tissues (Benham-Pyle et al., 2021), but these processes occur only at much later time points, i.e., a few days after injury, whereas activating wound responses in distal tissues is required within the first few hours.

We propose a model in which the distal wound responses act through providing feedback to the proximal responses at wounds (Figure 7). It is possible that planarian regeneration requires a secondary, activating signal provided by the distal tissues. Alternatively, uninjured tissues may produce inhibitory signals to block over-growth during homeostasis, but this inhibition is released when the wound signal arrives at distal tissues. Our data favor the second scenario. The existence of activating signals in distal wound responses would imply that we would have to supplement the specific cues in order to rescue the regeneration deficiencies caused by missing distal responses. This is, however, not what we observed. Instead, by amputating the distal tissues, which induces the proximal wound responses, is sufficient to rescue the regeneration. Identifying the molecular basis of this feedback loop will be an important avenue of future research. Altogether, our work highlights the functional crosstalk between spatially separated tissues not only during regeneration but may also be present in homeostatic tissues.

**Figure 7.**
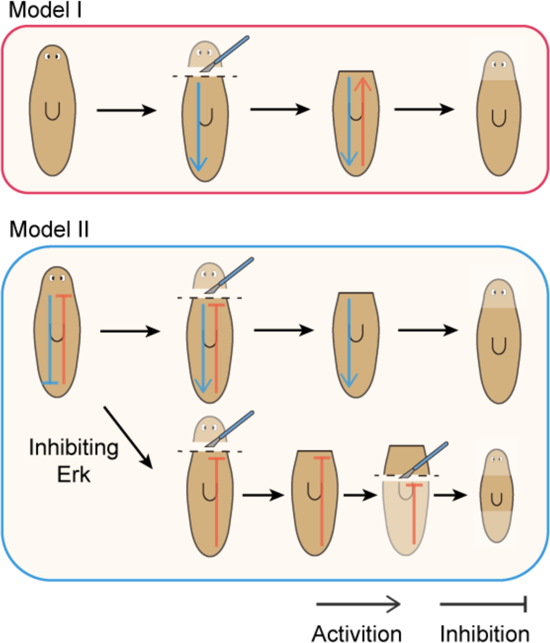
Models of feedback between proximal and distal tissues during regeneration. In Model I, after receiving the wound signal, the distal tissue sends an activating signal to the tissue at wounds to enable regeneration. In Model II, the inhibitory signal is removed after the distal tissue receives the wound signal so that regeneration can proceed. Blocking distal wound responses by inhibiting Erk propagation causes regeneration deficiencies, which can be rescued by a second injury removing the inhibitory signal.

## STAR Methods

### Key resources table

| REAGENT or RESOURCE | SOURCE | IDENTIFIER |
| --- | --- | --- |
| **Antibodies** |  |  |
| Anti-Digoxigenin-POD | Roche | 11207733910 |
| Anti-Digoxigenin-AP | Roche | 11093274910 |
| Anti-pErk | Santa Cruz Biotech | Sc-7383 |
| Anti-Erk1/2 | Cell Signaling | 9102L |
| Anti-actin | Hybridoma Bank | JLA20 |
| Goat anti-mouse IgG2a | LI-COR | 926-32351 |
| Goat anti-rabbit IgG | Biotium | 20067 |
| Goat anti-mouse IgM | Biotium | 20485 |
| Peroxidase goat anti-mouse IgG+IgM | Jackson ImmunoResearch | 115-035-044 |
| Anti-phospho-Histone H3 | Millipore Sigma | 04-817 |
| **Chemicals, peptides, and recombinant proteins** |  |  |
| TRIzol Reagent | Thermo Fisher Scientific | 15596026 |
| Dimethyl sulfoxide (DMSO) | Fisher Scientific | D1391 |
| U0126 | Cell Signaling | 9903S |
| Hoechst 33342 | Thermo Fisher Scientific | H3570 |
| Propidium iodide (PI) | Thermo Fisher Scientific | P3566 |
| DAPI | Sigma-Aldrich | D9542 |
| Dithiothreitol (DTT) | Thermo Fisher Scientific | R0861 |
| Sodium Dodecyl Sulfate (SDS) | Fisher Scientific | BP166 |
| cOmplete protease inhibitor | Millipore Sigma | 5892791001 |
| PhosSTOP | Millipore Sigma | 4906845001 |
| Benzonase | Millipore Sigma | E1014 |
| RNasin ribonuclease inhibitor | Promega | N2515 |
| FAST SYBR Green Master Mix | Bio-rad | 1725122 |
| Western blocking buffer | LI-COR | 927-60001 |
| Proteinase K solution | Thermo Fisher Scientific | 25-530-049 |
| Liberase TH | Millipore Sigma | 5401151001 |
| **Critical commercial assays** |  |  |
| RNeasy Mini Kit | Qiagen | 74104 |
| Universal Plus mRNA-seq Library Preparation Kit | TECAN | 0520-A01 |
| Chromium Single Cell v3.1 Library/Gel Bead Kit | 10x Genomics | 1000121 |
| **Deposited data** |  |  |
| RNAseq | GEO |  |
| Single cell RNAseq | GEO |  |
| **Oligonucleotides** |  |  |
| Forward and reverse primers for riboprobe and dsRNA synthesis | This paper, Table S3 | N/A |
| Vector pJC53.2 | (Collins et al., 2010) | Addgene 26536 |
| **Experimental models: organisms/strains** |  |  |
| Asexual *S. mediterranea* (strain CIW4) | (Newmark and Sánchez Alvarado, 2002) | N/A |
| Sexual S2F8b planarian strain | (Guo et al., 2017) | N/A |
| **Software and algorithms** |  |  |
| Bowtie2 | (Langmead and Salzberg, 2012) | N/A |
| DESeq2 | (Love et al., 2014) | N/A |
| UMI-tools | (Smith et al., 2017) | N/A |
| Cutadapt | (Martin, 2011) | N/A |
| BBKNN | (Polański et al., 2020) | N/A |
| SAM | (Tarashansky et al., 2019) | N/A |
| Fiji | N/A | http://fiji.sc/ |
| Python 3 | N/A | http://www.python.org |

### Experimental model and subject details

#### Animals

Asexual *S. mediterranea* (CIW4) were maintained in the dark at 20 °C in 0.5 g/L Instant Ocean Sea Salts supplemented with 0.1 g/L sodium bicarbonate. Sexual planarians were maintained in 0.75× Montjuïc salts. They were fed calf liver paste once or twice a week and starved for 7 days before all experiments. Asexual animals of ∼5 mm in length or sexuals of ∼1 cm in length were used for western blotting experiments.

## Method details

### Bulk RNAseq and data analysis

Total RNA was extracted using Trizol from five pooled planarian fragments for each sample. Three biological replicates for each condition were processed in parallel. Libraries were prepared using the Universal Plus mRNA-seq Library Preparation Kit (TECAN), and sequenced on an Illumina NextSeq platform. Reads were mapped to the dd_Smed_v6 transcriptome (http://planmine.mpi-cbg.de) (Rozanski et al., 2019) using bowtie2 with --sensitive flag (Langmead and Salzberg, 2012). The raw read counts from different isoforms of the same gene were lumped for downstream analysis. Pairwise differential expression analysis was performed using DESeq2 (Love et al., 2014). Heatmaps were generated using seaborn library with default parameters by calculating z-scores of normalized read counts generated by DESeq2.

#### Single-cell RNA sequencing and data analysis

Planarians were treated with either 0.25% DMSO or 25 µM U0126 in Instant Ocean for 24 h, and cut into three pieces 3 h before dissociation. To dissociate animals, planarians were finely minced with a razor blade. The minced tissues were then suspended in 3 mL of CMF (Ca/Mg-Free media: NaH_2_PO_4_ 480 mg/L, NaCl 960 mg/L, KCl 1.44 g/L, NaHCO_3_ 960 mg/L, HEPES 3.57 g/L, D-glucose 0.24 g/L, BSA 1 g/L, pH 7.4 in MillQ H_2_O) supplemented with equal volumes of DMSO or 25 µM U0126 (Cell Signaling) in DMSO for the control and Erk-inhibited groups respectively, and rocked for a total of 10 min, with gentle pipetting every 3 min. The cell suspension was serially filtered through 100, 70, 40, 30-µm mesh strainers to remove undissociated tissue chunks and cell aggregates. The filtered cell suspensions were centrifuged at 400 g for 5 min, and resuspended in 2 mL of CMF supplemented with DMSO or U0126. The cells were then incubated in Hoechst (10 µg/mL) for 30 min in the dark and propidium iodide (2 µg/mL) afterwards. Cells were sorted on a SONY SH800S based on live-dead gating into CMF containing 1% BSA.

Sorted cells were spun down at 500 g for 5 min and resuspended in CMF with 1% BSA to a final density of ∼1,000 cells per µL. Cells were then processed using 10x Genomics Chromium Controller and Chromium single cell v3.1 library/Gel Bead Kit. Amplified cDNA libraries were quantified using a bioanalyzer, and sequenced using Illumina Novaseq S4, generating in mean coverage of ∼ 34,000 and ∼31,000 read pairs per cell for the DMSO and U0126 treated cells, respectively. Sequenced reads were tagged with cell and molecular specific barcodes using UMI-tools (Smith et al., 2017), trimmed of primer and polyA sequences using cutadapt (Martin, 2011), and aligned to dd_Smed_v6 (Rozanski et al., 2019) using bowtie2 --sensitive parameter (Langmead and Salzberg, 2012).

Downstream preprocessing and analysis were performed using UMI counts, for which we lumped raw UMI counts from different isoforms of the same gene. Cells with fewer than 600 genes detected were filtered out, resulting in a final count of 13,276 and 14,793 cells for DMSO and U0126 treated samples. For each cell, we detected 2,252 genes and 6,483 UMI on average for the DMSO-treated sample and 2,275 genes and 6,043 UMI for the U0126-treated sample.

We normalized raw read counts for sequencing coverage such that each cell has a total read count equal to that of the median library size for all cells. The resulting counts were then added with a pseudo count of 1 and log-2 transformed. Integration of the control and Erk-inhibitor treated samples were performed using ridge regression on both technical effect and biological condition prior to applying BBKNN (Polański et al., 2020). 2D embedding was performed using the SAM algorithm (Tarashansky et al., 2019) with default parameters. Cells were annotated using cell-type specific markers from previously published RNAseq cell type atlas (Figure S2) (Fincher et al., 2018). All annotations are provided in **Table S1**. *p*-values on gene expression differences in cells treated with DMSO and U1026 were performed on each cell type separately via Mann-Whitney-Wilcoxon test in sciPy library with default parameters.

#### Western blotting

At specified time points after amputation, animals were incubated in zinc fixative (100 nM ZnCl_2_ in ethanol) for 45 min at 4 °C, then cut into four pieces along anterior-posterior axis, each measuring ∼1 mm in length, on cool packs (Figure S4B). The most anterior tissue piece (P1) around the wound and the distal tissue piece (P3) between the pharynx and tail were used in U0126-treatment and *myoD* RNAi experiments because they exhibited largest dynamic ranges of Erk activity post amputation (Figure 2B). From the same proximal-distal position, 5-10 tissue pieces were pooled and lysed by mechanical pestle homogenization in urea lysis buffer (6 M urea, 2% SDS, 130 mM DTT, 3.5 U/ml Benzonase, 1x protease inhibitor cocktail, 1x phosphatase inhibitor) per experiment as described before (Stückemann et al., 2017). Protein concentrations were determined by a Nanodrop spectrophotometer based on absorbance at 280 nm. Western blotting was performed following an established protocol (Stückemann et al., 2017). Anti-pErk (Santa Cruz Biotech, sc-7383, 1:1000), anti-Erk (Cell Signaling, 9102L, 1:250), and anti-actin (JLA20, Hybridoma Bank, 1:7500) were used as primary antibodies. The specificity of these Erk antibodies was validated by U0126 treatment and *erk* RNAi (Figure S4A).

Goat anti-mouse IgG2a (LI-COR, 926-32351, 1:10000), Goat anti-rabbit IgG (Biotium, 20067, 1:20000), and Goat anti-mouse IgM (Biotium, 20485, 1:10000) were used as secondary antibodies. Membranes were imaged on a LI-COR Odyssey imager. The images were quantified using ImageJ Fiji software. Erk activation was measured by the ratio between pErk and total Erk and normalized to the 0 hpa activity in each group. The peak time of Erk activation was determined by fitting the activation vs. time with polynomials.

#### Cloning, in situ hybridization and immunostaining

Gene fragments were amplified from cDNA using oligonucleotide primers listed in **Table S3** and cloned into the vector pJC53.2 (Addgene Plasmid ID: 26536) (Collins et al., 2010). RNA probes were synthesized through *in vitro* transcription as described previously (King and Newmark, 2013). Whole-mount in situ hybridization (WISH) and immunostaining were performed following the established protocols (King and Newmark, 2013).

Anti-phospho-Histone H3 (Millipore Sigma, 04-817, 1:300) was used for H3P staining. For single muscle cell fluorescence in situ hybridization (FISH)/immunostaining, animals were amputated to induce Erk activation and then dissociated at 3 hpa in CMF supplemented with a 1:10 dilution of Liberase TH (Millipore Sigma, 5401135001) into single cells (Witchley et al., 2013) and processed as described recently (Grohme et al., 2021).

#### RNAi and drug treatment

dsRNA was prepared from *in vitro* transcription reactions using PCR-generated templates with flanking T7 promoters, followed by ethanol precipitation, and annealed after resuspension as described before (Collins et al., 2010). dsRNA was mixed with liver paste at a concentration of 100 ng/μl and fed to animals every 4 d for 10 times. The RNAi animals were starved for 7 d before experiments. In all RNAi experiments, dsRNA matching the *ccdB* and *camR*-containing insert of pJC53.2 (Collins et al., 2010) was used as the negative control.

The irreversible Mek inhibitor, U0126, was first dissolved in DMSO to 10 mM and diluted in Instant Ocean to desired final concentrations. Animals were treated for 4 h before amputations, and changed fresh Instant Ocean containing U0126 daily throughout regeneration.

#### Mathematical modeling

We adopted a recently developed diffusive signaling relay model (Dieterle et al., 2020) to investigate the effects of cell length, density, orientation, and diversity on Erk activity propagation. This model considers a relay in which cells sense the local concentration of a diffusive signaling molecule and participate in the release of the same molecule to activate neighboring cells. Each cell occupies a specified space in a 2D plane and activates when the average concentration of activator within its occupied space exceeds a certain threshold (Figure S5A-C).

For cells around the wound, the concentration of the signaling molecule *c*(*Z, t*) can be written as where

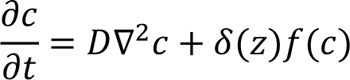

*D* is the diffusion coefficient of the signaling molecule, the Dirac function *δ*(*Z*) accounts for the fact that the cells respond to local activator concentrations, and *f*(*c*) is the source function accounting for the rate at which the signaling molecule is produced at the wound site. Similarly, for relay cells that are away from wounds:

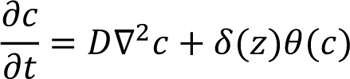

where *θ*(*c*) is the activation function that can be described as a step function with threshold *C*_*th*_. When the local signaling molecule concentration exceeds *C*_*th*_, the cell is activated and starts to produce the activator. The assumption that the rate of production of *c* depends on the presence of *c* above the threshold provides the system with the positive feedback required for trigger waves. For receiving cells that only receives the signal without amplifying it:

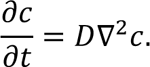

To simulate wave propagation, *D* was varied around 0.1 µm^2^/s based on previously inferred diffusion coefficients of intercellular Erk activator (Hayden et al., 2021; De Simone et al., 2021). This slow diffusion is also justified by the narrow Erk wave width (< 1 mm) observed in our system as fast diffusion would significantly broaden the wave front. Cell width was defaulted to be 10 μm and length was fixed at 10 μm for small cells but varied between 10-100 μm for long cells. Density of long cells was defined by the fraction of the 2D plane occupied by long cells. Orientation factor of long cells was defined by the fraction of long cells extending along the direction of wave propagation whereas other long cells are perpendicular to the wave direction.

For numerical simulations, source function and activation function were set to be:

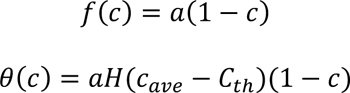

where *a* = 12 *S*^-1^, *H* is the Heaviside step function, *c*_ave_ is the average concentration of cell region, and *C*_*th*_ = 0.1. All simulations were performed on a 1 mm by 0.2 mm plane with 10 μm grid size. The simulation time step was 0.01 min. *c* was set to be 0 everywhere on the plane as the initial condition. Our simulation validated that this model can capture the essential dynamics of trigger waves (Figure S5B-C). Signal propagation speed was calculated through the mean displacement of the wave front, position at which average signal activity exceeds 0.1 (Figure 56C), per unit time.

## Acknowledgements.

We thank LE O’Brien, S Granick, S Di Talia, M Wu, and members of Wang lab and Ferrell lab for critical discussions and J Gibson for assistance in cell sorting. YF and XZ are Bio-X Stanford Interdisciplinary Graduate Fellows. CC is supported by a NSF Graduate Research Fellowship and a Stanford Graduate Fellowship. BW is a Beckman Young Investigator. This work is supported by NIH grant 1R35GM138061 to BW and 5R35GM131792 to JF.

## Authors contributions

Conceptualization: YF, JF, and BW; Methodology: YF, CC, PL, and XZ; Investigation: YF, CC, PL, and XZ; Formal analysis: YF and CC; Validation: YF; Writing: YF, JF, and BW with feedback from all other authors; Funding acquisition: BW; Supervision: BW.

## Supplemental figures

**Figure S1.**
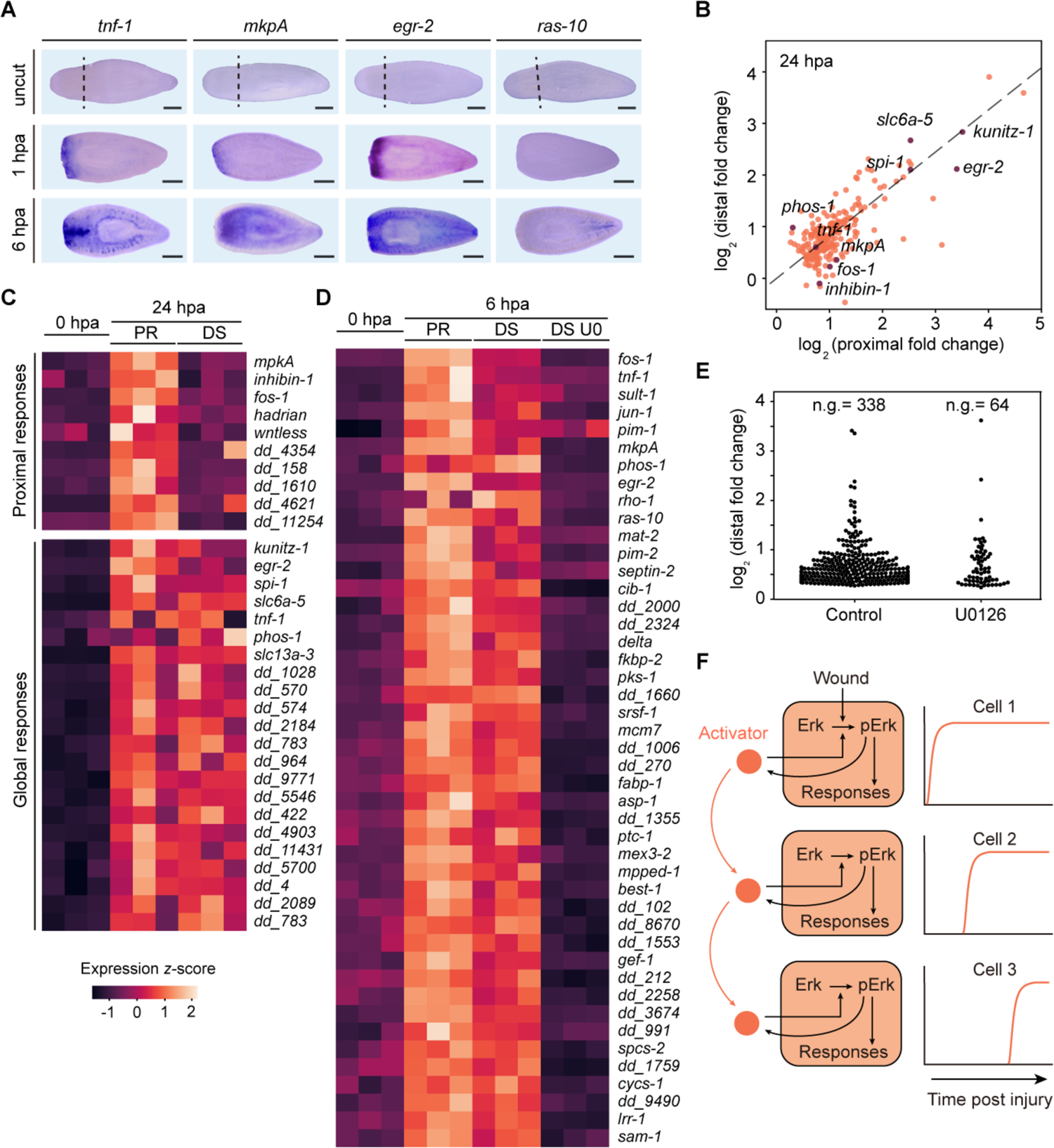
Systemic wound responses in planarian regeneration, related to Figure 1. (A) WISH images showing activation of wound responses first in proximal and then in distal tissues. *tnf-1, mkpA*, and *egr-2* are only activated around the injury site at 1 hpa, and their expressions spread to the rest of the body within 6 hpa. Induction of *ras-10* expression is specific in the tail but only becomes activated at 6 hpa. Dashed lines: amputation plane. Scale bars: 1 mm. (B) Comparison of gene upregulation at 24 hpa in proximal and distal tissues. Fold changes were calculated from three biological replicates. Each gene plotted has p-value<0.001 (two-sided Welch’s t-test) and log_2_(fold change)>0.5 in either proximal or distal group. Note that the top upregulated genes are different from those observed at 6 hpa. (C) Heatmap of upregulated example wound response genes at 24 hpa. (D) Heatmap of globally upregulated wound response genes at 6 hpa. U0126 treatment blocked the upregulation. In (C-D), PR, proximal; DS, distal; U0, 25 μM U0126 treated. Data from three biological replicates are shown. (E) Fold changes of wound response genes at 6 hpa (p-value<0.001, two-sided Welch’s t-test, compared to 0 hpa) in distal tissues. U0126 treatment (25 μM) blocks the upregulation of ∼80% of wound response genes in distal tissues; n.g., number of genes. (F) Schematic of Erk activity propagation. Activator (magenta) induces Erk phosphorylation. Activated Erk triggers transcriptional activation of wound responses and induces the release of more activators to extracellular space, which can induce a positive feedback in the same cell, or diffuse to cells adjacent in space and trigger Erk activation therein. This signaling mechanism can result in Erk activity to be relayed from cell to cell with constant speed and amplitude. Common Erk activating pathways include Egf (Gagliardi et al., 2021; Kiyatkin et al., 2020) and Fgf (De Simone et al., 2021) pathways. However, the planarian genome lacks conserved Egf and Fgf homologs based on BLAST search. Egf and Fgf receptors are present in the planarian genome and have been shown to play complex roles in regulating stem cell fates (Lei et al., 2016) and body plan formation (Cebrià et al., 2002; Scimone et al., 2016; Umesono et al., 2013). Therefore, it is difficult to dissect the direct link between Egf/Fgf and Erk pathways.

**Figure S2.**
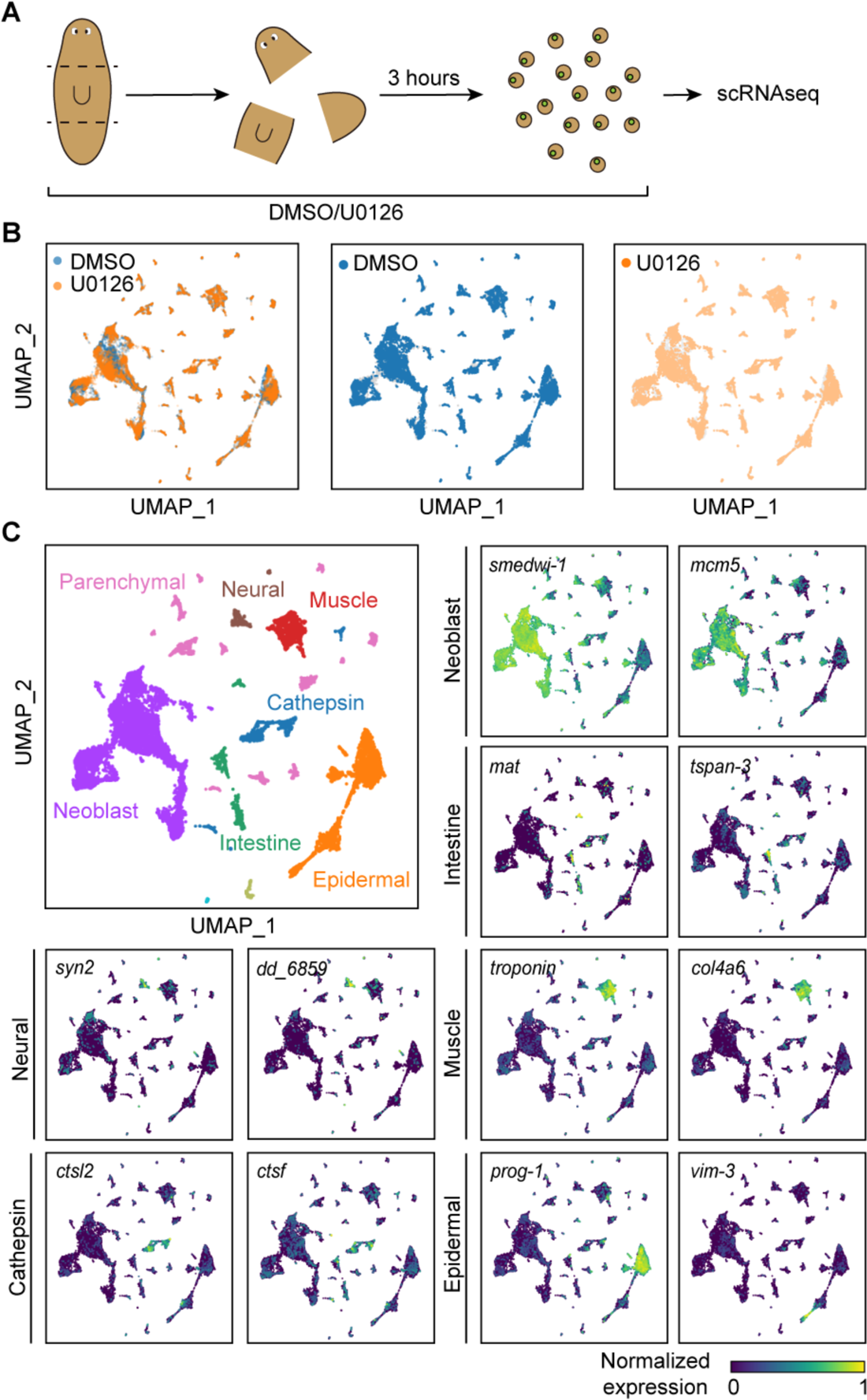
Single-cell RNAseq to capture cell type-specific Erk-dependent wound responses, related to Figure 1. (A) Schematic showing the strategy to measure cell type-specific wound responses using scRNAseq. (B) All cell clusters contain cells from both conditions: DMSO-treated (blue, 13,276 cells) and 25 μM U0125-treated (orange, 14,793 cells). (C) Annotation of major cell types and example marker gene expression patterns. Neoblast: *smedwi-1* and *mcm5*; intestine: *mat* and *tspan-3*; muscle: *troponin* and *col4a6*; cathepsin: *ctsl2* and *ctsf*; epidermal: *prog-1* and *vim-2*; neural: *dd_6859* and *syn2*.

**Figure S3.**
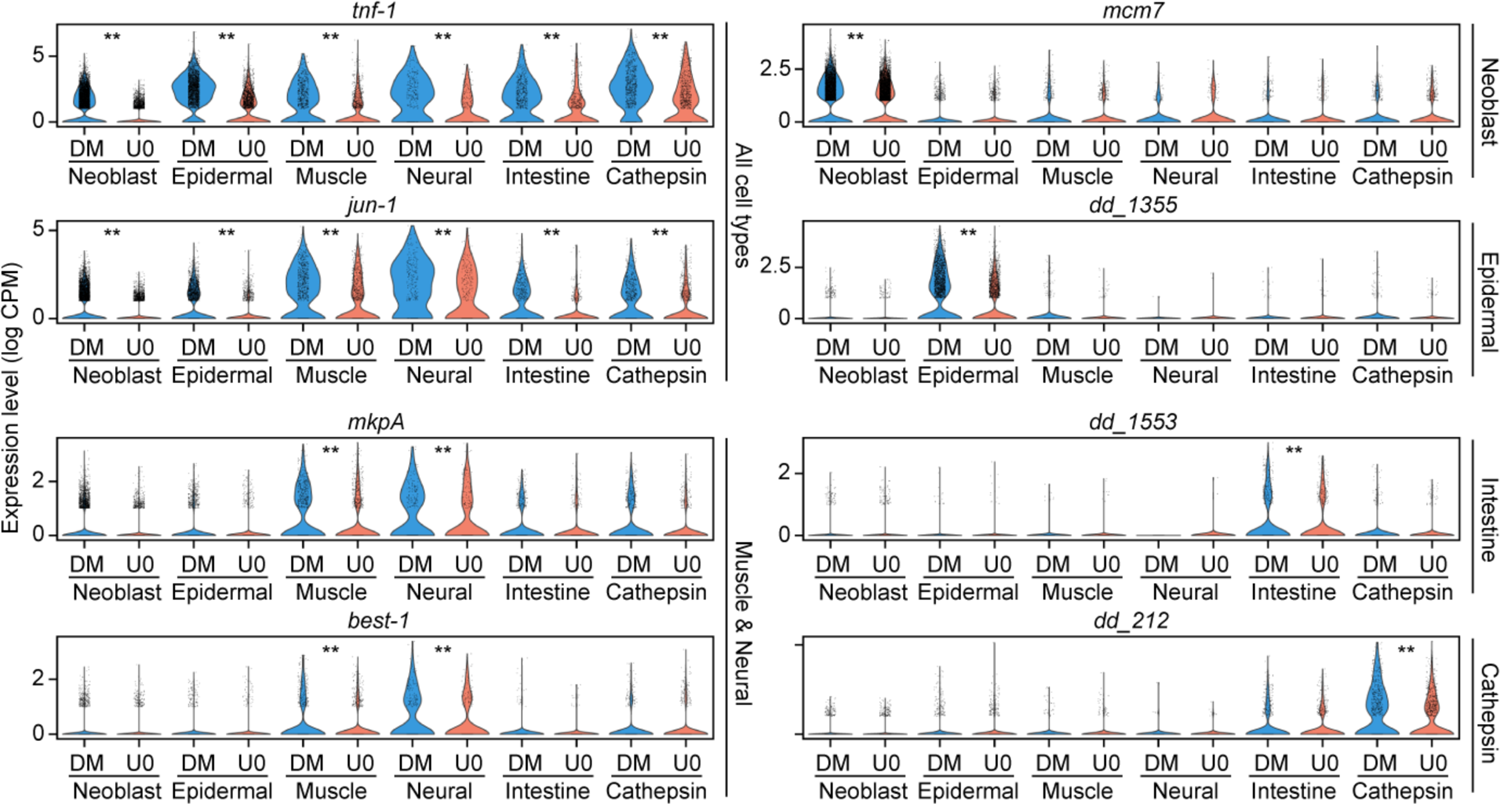
scRNAseq analysis identifies cell type-specific Erk-dependent wound response genes, related to Figure 1 Violin plots showing the expression distribution of select wound response genes in animals treated by DMSO (DM) or 25 μM U0126 (U0). Points: data of individual cells. **, p<0.01 between DMSO and 25 μM U0126 treated groups in specific cell types, two-sided Mann-Whitney-Wilcoxon test.

**Figure S4.**
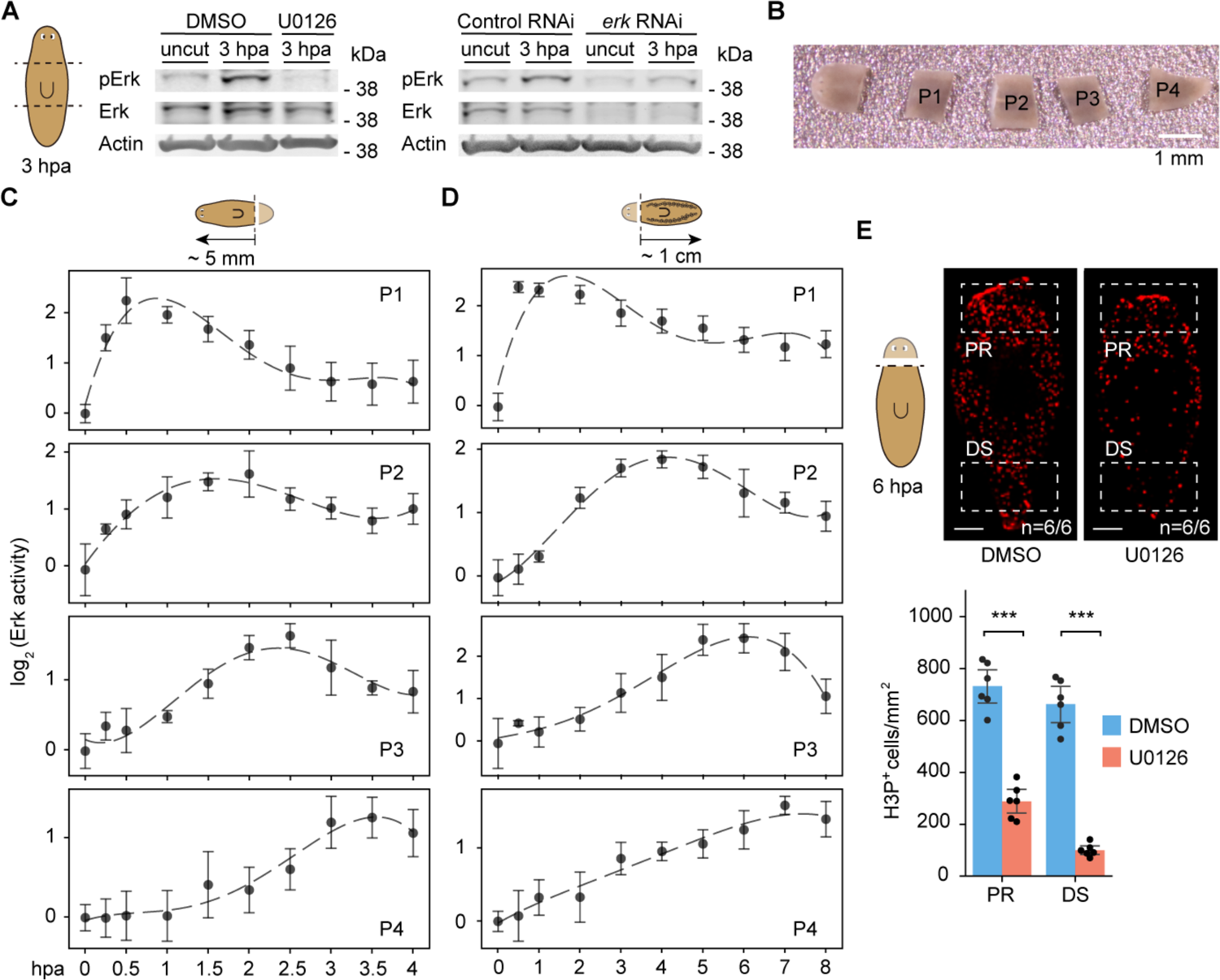
Measuring Erk activity using western blotting, related to Figure 2 and 3 (A) Western blot images showing that 25 μM U0126 treatment eliminates pErk signal (left) and that *erk* RNAi decreases total Erk signal (right). These two experiments validate the specificity of the antibodies. (B) A representative image showing that fixed worms are cut into pieces for protein extraction to detect spatiotemporal patterns of Erk activity. (C-D) Erk activity after posterior amputation (C) and in a sexual biotype with a longer body plan (D). Dashed lines: polynomial fit. (E) (Top) Anti-H3P labels mitotic cells at 6 hpa in animals treated with DMSO or 25 μM U0126. Dashed line: amputation plane. (Bottom) Number of H3P^+^ cells counted in the proximal (PR) and distal (DS) regions (boxes in images). ∗∗∗, p < 0.001, two-sided Welch’s t-test; error bars: 95% confidence interval (CI). Scale bars: 1 mm in (B), 500 μm in (E).

**Figure S5.**
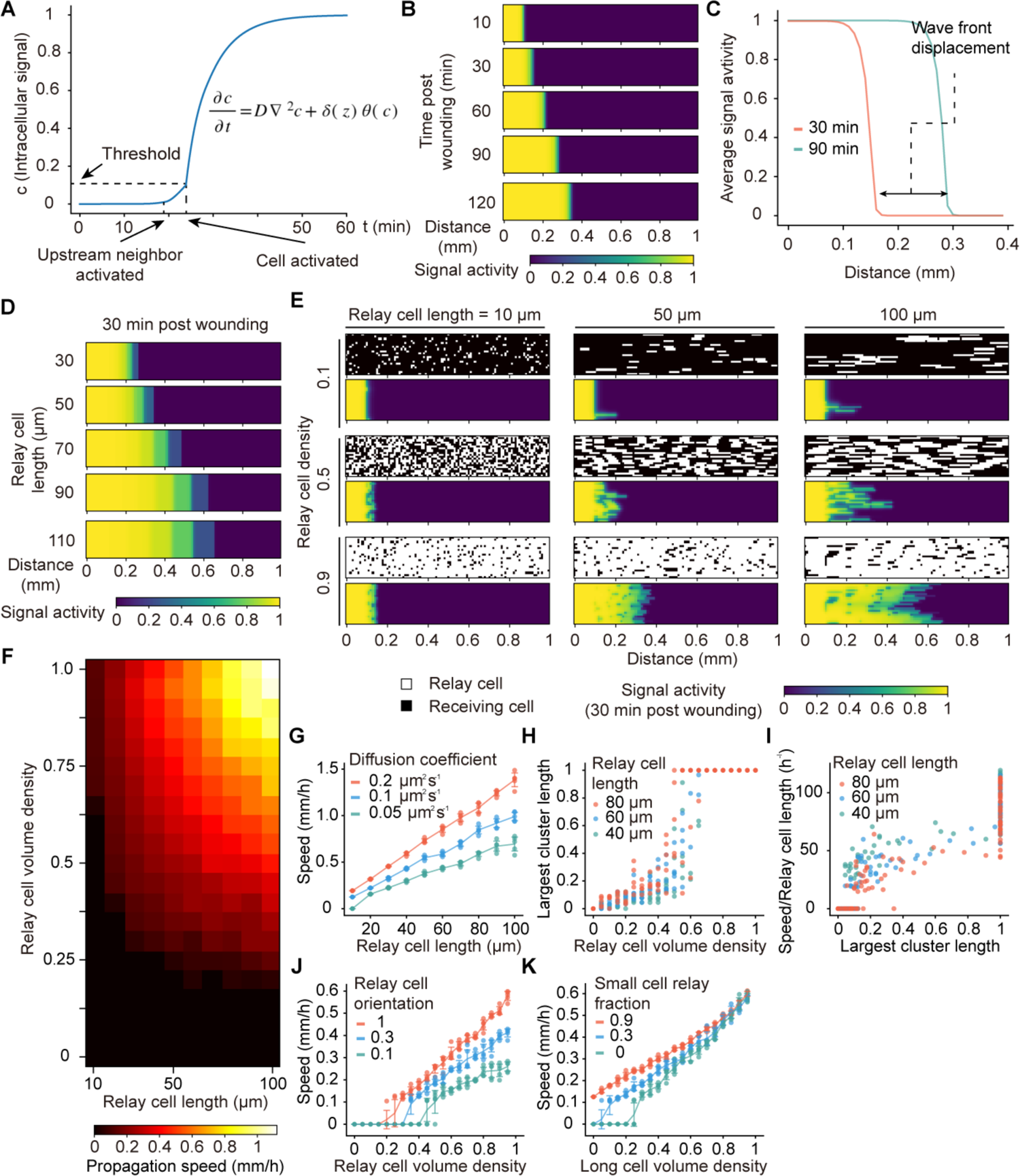
A diffusive signaling relay model to simulate biochemical activity propagation in heterogeneous tissues, related to Figure 5 and 6 (A) Single cell signaling activation dynamics during wave propagation. The time of activation (when signaling molecule concentration exceeds the threshold, *C*_*th*_) for the cell and its upstream neighbor is indicated by arrows. (B) Simulated wave propagation. Under this condition, the wave propagates at a constant speed (∼150 μm/h). (C) Mean signal activity vs. position. Wave fronts are defined at the position where mean signal activity exceeds 0.1. The displacement of wave fronts per unit time is used to calculate wave speed. (D) Erk activity propagates faster through longer cells. These simulations only contain uniform populations of long cells. (E) Spatial distributions of signal relay cells (top) and signal activity (bottom) in heterogeneous tissues with varying long cell lengths and volume densities. Note that cells that do not relay signal also can be activated in this model. (F) Heatmap showing the speed of signal propagation increases with relay cell length and volume density. (G) The trend that signal propagation speed increases with relay cell length is robust to varying ligand diffusion coefficient. Long cell volume density is fixed at 0.9. (H) The length of the largest long cell cluster along the direction of wave propagation, normalized by the simulation box length, vs. relay cell volume density. Length of 1 indicates that the largest cluster spans the entire system. (I) Replotting the data in (H) to show that the signal propagation speed increases with the length of the largest long cell cluster. The speed is in the unit of long cell length. (J) Signal propagation speed increases with relay cell density in simulations varying long cell orientation factors. The discontinuity in the speed is observed at the percolation transition and shifts towards higher volume density with smaller long cell orientation factor. (K) Signal propagation speed increases with long cell volume density in simulations varying fractions of small cells that can relay the signal. Increasing fraction of relaying small cells shifts the percolation transition towards lower long cell density. In all simulations shown in (E-J), small cells only can receive but cannot relay signal. Orientation factor is fixed at 1 except for (J). In (J-K), long cells are set to be 50 μm in length.

**Figure S6.**
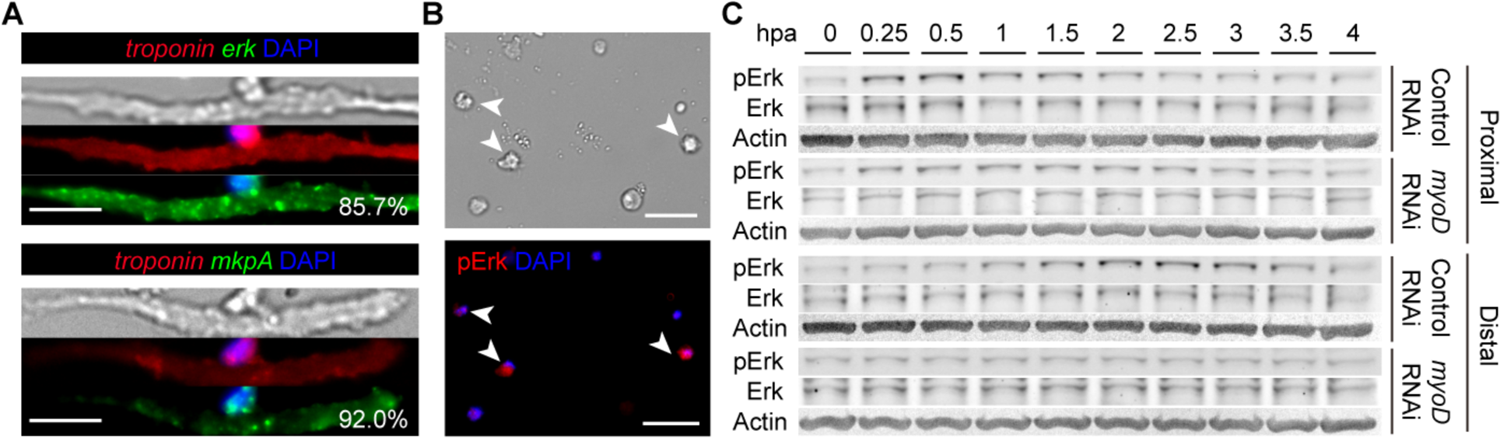
Muscles are essential for Erk activity propagation, related to Figure 5 (A) Bright field and FISH images of isolated muscle cells (*troponin^+^*) expressing *erk* (top) and *mkpA* (bottom), a conserved Erk-responsive dual-specificity phosphatases (Tasaki et al., 2011; Umesono et al., 2013). %, *erk*^+^ and *mkp*A^+^ fractions of muscle cells. (B) Bright field and pErk immunostaining images of isolated cells to show cell types other than muscles can activate Erk pathway as well. Arrows: pErk^+^ cells. In (A-B), scale bars: 20 μm. (C) Representative western blot images showing Erk is not activated in tissues distal to wound after *myoD* RNAi. Actin was used as loading control.

## Supplemental Tables

**Table S1.** Single-cell annotations with batch information, number of genes detected per cell, and number of UMI detected per cell, related to Figure 1.

**Table S2.** Contig numbers for all mentioned genes, related to Figure 1 and 5.

**Table S3.** Primers used for cloning, related to the STAR Methods.

## References

Aoki, K., Kondo, Y., Naoki, H., Hiratsuka, T., Itoh, R.E., and Matsuda, M. (2017). Propagating wave of ERK activation orients collective cell migration. Dev. Cell 43, 305–317.e5.

Benham-Pyle, B.W., Brewster, C.E., Kent, A.M., Mann, F.G., Chen, S., Scott, A.R., Box, A.C., and Sánchez Alvarado, A. (2021). Identification of rare, transient post-mitotic cell states that are induced by injury and required for whole-body regeneration in Schmidtea mediterranea. Nat. Cell Biol. 23, 939–952.

Cebrià, F., Kobayashi, C., Umesono, Y., Nakazawa, M., Mineta, K., Ikeo, K., Gojobori, T., Itoh, M., Taira, M., Sánchez Alvarado, A., et al. (2002). FGFR-related gene nou-darake restricts brain tissues to the head region of planarians. Nature 419, 620–624.

Chang, J.B., and Ferrell, J.E. (2013). Mitotic trigger waves and the spatial coordination of the Xenopus cell cycle. Nature 500, 603–607.

Cheng, X., and Ferrell, J.E. (2018). Apoptosis propagates through the cytoplasm as trigger waves. Science 361, 607–612.

Chera, S., Ghila, L., Dobretz, K., Wenger, Y., Bauer, C., Buzgariu, W., Martinou, J.C., and Galliot, B. (2009). Apoptotic cells provide an unexpected source of Wnt3 signaling to drive *Hydra* head regeneration. Dev. Cell 17, 279–289.

Collins, J.J., Hou, X., Romanova, E. V., Lambrus, B.G., Miller, C.M., Saberi, A., Sweedler, J. V., and Newmark, P.A. (2010). Genome-wide analyses reveal a role for peptide hormones in planarian germline development. PLoS Biol 8, e1000509.

Deneke, V.E., and Di Talia, S. (2018). Chemical waves in cell and developmental biology. J. Cell Biol. 217, 1193–1204.

Dieterle, P.B., Min, J., Irimia, D., and Amir, A. (2020). Dynamics of diffusive cell signaling relays. eLife 9, e61771.

DuBuc, T.Q., Traylor-Knowles, N., and Martindale, M.Q. (2014). Initiating a regenerative response; cellular and molecular features of wound healing in the cnidarian *Nematostella vectensis*. BMC Biol. 12, 24.

Fincher, C.T., Wurtzel, O., de Hoog, T., Kravarik, K.M., and Reddien, P.W. (2018). Cell type transcriptome atlas for the planarian *Schmidtea mediterranea*. Science 360, eaaq1736.

Gagliardi, P.A., Dobrzyński, M., Jacques, M.A., Dessauges, C., Ender, P., Blum, Y., Hughes, R.M., Cohen, A.R., and Pertz, O. (2021). Collective ERK/Akt activity waves orchestrate epithelial homeostasis by driving apoptosis-induced survival. Dev. Cell 56, 1712–1726.e6.

Gaviño, M.A., Wenemoser, D., Wang, I.E., and Reddien, P.W. (2013). Tissue absence initiates regeneration through Follistatin-mediated inhibition of Activin signaling. eLife 2, e00247.

Gelens, L., Anderson, G.A., and Ferrell, J.E. (2014). Spatial trigger waves: positive feedback gets you a long way. Mol. Biol. Cell 25, 3486–3493.

Grohme, M., Frank, O., and Rink, J. (2021). Preparing planarian cells for high-content fluorescence microscopy using RNA in situ hybridization and immunocytochemistry. Preprints 2021110256.

Guo, L., Zhang, S., Rubinstein, B., Ross, E., and Sánchez Alvarado, A. (2017). Widespread maintenance of genome heterozygosity in Schmidtea mediterranea. Nat. Ecol. Evol 1, 0019.

Halme, A., Cheng, M., and Hariharan, I.K. (2010). Retinoids regulate a developmental checkpoint for tissue regeneration in Drosophila. Curr Biol 20, 458–463.

Hasegawa, T., Nakajima, T., Ishida, T., Kudo, A., and Kawakami, A. (2015). A diffusible signal derived from hematopoietic cells supports the survival and proliferation of regenerative cells during zebrafish fin fold regeneration. Dev. Biol. 399, 80–90.

Hayden, L.D., Poss, K.D., De Simone, A., and Di Talia, S. (2021). Mathematical modeling of Erk activity waves in regenerating zebrafish scales. Biophys. J. 120, 4287–4297.

Hiratsuka, T., Fujita, Y., Naoki, H., Aoki, K., Kamioka, Y., and Matsuda, M. (2015). Intercellular propagation of extracellular signal-regulated kinase activation revealed by *in vivo* imaging of mouse skin. eLife 4, e05178.

Hirose, K., Payumo, A.Y., Cutie, S., Hoang, A., Zhang, H., Guyot, R., Lunn, D., Bigley, R.B., Yu, H., Wang, J., et al. (2019). Evidence for hormonal control of heart regenerative capacity during endothermy acquisition. Science 364, 184–188.

Johnson, K., Bateman, J., Di Tommaso, T., Wong, A.Y., and Whited, J.L. (2018). Systemic cell cycle activation is induced following complex tissue injury in axolotl. Dev. Biol. 433, 461–472.

Khariton, M., Kong, X., Qin, J., and Wang, B. (2020). Chromatic neuronal jamming in a primitive brain. Nat. Phys. 16, 553–557.

Kikuchi, K., Holdway, J.E., Major, R.J., Blum, N., Dahn, R.D., Begemann, G., and Poss, K.D. (2011). Retinoic acid production by endocardium and epicardium is an injury response essential for zebrafish heart regeneration. Dev. Cell 20, 397–404.

King, R.S., and Newmark, P.A. (2013). In situ hybridization protocol for enhanced detection of gene expression in the planarian *Schmidtea mediterranea*. BMC Dev. Biol. 13, 8.

Kiyatkin, A., van Alderwerelt van Rosenburgh, I.K., Klein, D.E., and Lemmon, M.A. (2020). Kinetics of receptor tyrosine kinase activation define ERK signaling dynamics. Sci. Signal. 13, eaaz5267.

Langmead, B., and Salzberg, S.L. (2012). Fast gapped-read alignment with Bowtie 2. Nat. Methods 9, 357–359.

Lei, K., Thi-Kim Vu, H., Mohan, R.D., McKinney, S.A., Seidel, C.W., Alexander, R., Gotting, K., Workman, J.L., and Sánchez Alvarado, A. (2016). Egf signaling directs neoblast repopulation by regulating asymmetric cell division in planarians. Dev. Cell 38, 413–429.

Lim, Y., Khariton, M., Lane, K., Shiver, A., Ng, K., Bray, S., Qin, J., Huang, K.C., and Wang, B. (2019). Mechanically resolved imaging of bacteria using expansion microscopy. PLoS Biol. 17, e3000268.

Losner, J., Courtemanche, K., and Whited, J.L. (2021). A cross-species analysis of systemic mediators of repair and complex tissue regeneration. Npj Regen. Med. 6, 21.

Love, M.I., Huber, W., and Anders, S. (2014). Moderated estimation of fold change and dispersion for RNA-seq data with DESeq2. Genome Biol. 15, 550.

Mace, K.A., Pearson, J.C., and McGinnis, W. (2005). An epidermal barrier wound repair pathway in Drosophila is mediated by grainy head. Science 308, 381–385.

Martin, M. (2011). Cutadapt removes adapter sequences from high-throughput sequencing reads. EMBnet.Journal 17, 10–12.

Müller, P., Rogers, K.W., Yu, S.R., Brand, M., and Schier, A.F. (2013). Morphogen transport. Development 140, 1621–1638.

Newmark, P.A., and Sánchez Alvarado, A. (2002). Not your father’s planarian: A classic model enters the era of functional genomics. Nat. Rev. Genet. 3, 210–219.

Owlarn, S., Klenner, F., Schmidt, D., Rabert, F., Tomasso, A., Reuter, H., Mulaw, M.A., Moritz, S., Gentile, L., Weidinger, G., et al. (2017). Generic wound signals initiate regeneration in missing-tissue contexts. Nat. Commun. 8, 2282.

Payzin-Dogru, D., Wilson, S.E., Blair, S.J., Erdogan, B., Hossain, S., Cammarata, L., Velazquez Matos, J., and Whited, J.L. (2021). Nerve-mediated amputation-induced stem cell activation primes distant appendages for future regeneration events in axolotl. Https://Biorxiv.Org/Content/10.1101/2021.12.29.474455v1.

Pellettieri, J., Fitzgerald, P., Watanabe, S., Mancuso, J., Green, D.R., and Sánchez Alvarado, A. (2010). Cell death and tissue remodeling in planarian regeneration. Dev. Biol. 338, 76–85.

Petersen, C.P., and Reddien, P.W. (2011). Polarized notum activation at wounds inhibits Wnt function to promote planarian head regeneration. Science 332, 852–855.

Polański, K., Young, M.D., Miao, Z., Meyer, K.B., Teichmann, S.A., and Park, J.E. (2020). BBKNN: Fast batch alignment of single cell transcriptomes. Bioinformatics 36, 964–965.

Ricci, L., and Srivastava, M. (2018). Wound-induced cell proliferation during animal regeneration. WIREs Dev. Biol. 7, e321.

Rink, J.C. (2013). Stem cell systems and regeneration in planaria. Dev. Genes Evol. 223, 67– 84.

Roberts-Galbraith, R.H., and Newmark, P.A. (2013). Follistatin antagonizes Activin signaling and acts with Notum to direct planarian head regeneration. Proc. Natl. Acad. Sci. U S A 110, 1363–1368.

Rodgers, J.T., King, K.Y., Brett, J.O., Cromie, M.J., Charville, G.W., Maguire, K.K., Brunson, C., Mastey, N., Liu, L., Tsai, C.R., et al. (2014). MTORC1 controls the adaptive transition of quiescent stem cells from G0 to GAlert. Nature 510, 393–396.

Rodgers, J.T., Schroeder, M.D., Ma, C., and Rando, T.A. (2017). HGFA is an injury-regulated systemic factor that induces the transition of stem cells into GAlert. Cell Rep. 19, 479–486.

Rozanski, A., Moon, H., Brandl, H., Martín-Durán, J.M., Grohme, M.A., Hüttner, K., Bartscherer, K., Henry, I., and Rink, J.C. (2019). PlanMine 3.0—improvements to a mineable resource of flatworm biology and biodiversity. Nucleic Acids Res. 47, D812–D820.

Scimone, M.L., Cote, L.E., Rogers, T., and Reddien, P.W. (2016). Two FGFRL-Wnt circuits organize the planarian anteroposterior axis. ELife 5, e12845.

Scimone, M.L., Cote, L.E., and Reddien, P.W. (2017). Orthogonal muscle fibres have different instructive roles in planarian regeneration. Nature 551, 623–628.

De Simone, A., Evanitsky, M.N., Hayden, L., Cox, B.D., Wang, J., Tornini, V.A., Ou, J., Chao, A., Poss, K.D., and Di Talia, S. (2021). Control of osteoblast regeneration by a train of Erk activity waves. Nature 590, 129–133.

Smith, T., Heger, A., and Sudbery, I. (2017). UMI-tools: Modeling sequencing errors in Unique Molecular Identifiers to improve quantification accuracy. Genome Res. 27, 491–499.

Srivastava, M. (2021). Beyond casual resemblance: Rigorous frameworks for comparing regeneration across species. Annu. Rev. Cell Dev. Biol. 37, 415–440.

Stauffer, D., and Aharony, A. (1994). Introduction to Percolation Theory (Taylor & Francis).

Stückemann, T., Cleland, J.P., Werner, S., Thi-Kim Vu, H., Bayersdorf, R., Liu, S.Y., Friedrich, B., Jülicher, F., and Rink, J.C. (2017). Antagonistic self-organizing patterning systems control maintenance and regeneration of the anteroposterior axis in planarians. Dev. Cell 40, 248–263.e4.

Sun, F., Ou, J., Shoffner, A., Luan, Y., Yang, H., Song, L., Safi, A., Cao, J., Yue, F., Crawford, G., et al. (2022). Enhancer selection dictates gene expression responses in remote organs during tissue regeneration. Nat. Cell Biol. 24, 685–696.

Tarashansky, A.J., Xue, Y., Li, P., Quake, S.R., and Wang, B. (2019). Self-assembling manifolds in single-cell RNA sequencing data. eLife 8, e48994.

Tasaki, J., Shibata, N., Nishimura, O., Itomi, K., Tabata, Y., Son, F., Suzuki, N., Araki, R., Abe, M., Agata, K., et al. (2011). ERK signaling controls blastema cell differentiation during planarian regeneration. Development 138, 2417–2427.

Tewari, A.G., Stern, S.R., Oderberg, I.M., and Reddien, P.W. (2018). Cellular and molecular responses unique to major injury are dispensable for planarian regeneration. Cell Rep. 25, 2577–2590.

Tursch, A., Bartsch, N., and Holstein, T. (2020). MAPK signaling links the injury response to Wnt-regulated patterning in Hydra regeneration. Https://Biorxiv.Org/Content/10.1101/2020.07.06.189795v1.

Tyson, J.J., and Keener, J.P. (1988). Singular perturbation theory of traveling waves in excitable media (a review). Physica D 32, 327–361.

Umesono, Y., Tasaki, J., Nishimura, Y., Hrouda, M., Kawaguchi, E., Yazawa, S., Nishimura, O., Hosoda, K., Inoue, T., and Agata, K. (2013). The molecular logic for planarian regeneration along the anterior-posterior axis. Nature 500, 73–76.

Wenemoser, D., and Reddien, P.W. (2010). Planarian regeneration involves distinct stem cell responses to wounds and tissue absence. Dev. Biol. 344, 979–991.

Wenemoser, D., Lapan, S.W., Wilkinson, A.W., Bell, G.W., and Reddien, P.W. (2012). A molecular wound response program associated with regeneration initiation in planarians. Genes & Dev. 26, 988–1002.

Winfree, A.T. (1974). Two kinds of wave in an oscillating chemical solution. Faraday Symp. Chem. Soc. 9, 38–46.

Witchley, J.N., Mayer, M., Wagner, D.E., Owen, J.H., and Reddien, P.W. (2013). Muscle cells provide instructions for planarian regeneration. Cell Rep. 4, 633–641.

